# Squash flowers as microhabitats: the effects of floral temperature and humidity on pollen viability and visitor behavior

**DOI:** 10.1101/2025.02.24.639912

**Authors:** Ajinkya Dahake, Charlotte Coates, Diana Obregon, Jonathan Chai, Patricia Nunes-Silva, Peter Kevan, Robert A. Raguso

## Abstract

Flowers represent intricate microcosms shaped by their chemical and micrometeorological properties. Notable examples include thermogenic flowers that create a warm microclimate for visitors and residents. We document a distinct microclimate within the large flowers of both wild and domesticated squashes (*Cucurbita spp*.). Unlike thermogenic flowers, squash flower temperatures remain near ambient, but their humidity consistently exceeds ambient levels from bud to senescence, resulting from stomatal and petal transpiration rather than nectar evaporation. Experimentally reducing humidity in greenhouse-grown male squash flowers results in significant pollen tube rupture, directly impacting plant fitness. To explore the role of floral humidity within a broader ecological context, we performed similar humidity manipulations on squash farms to assess impacts on the behavior of their specialist squash bee pollinator (*Xenoglossa pruinosa*), generalist pollinators (bumblebees, honeybees), and specialist herbivores (cucurbit beetles). Experimentally reduced floral humidity lower visitation frequency by squash bees but have no effect on generalist pollinators. Manipulation of floral humidity did not influence the foraging duration of any pollinators but impacted the residence of male squash bees in wilted flowers compared with unmanipulated flowers. Finally, there was a positive correlation between the dryness of the ambient air and the abundance of squash bees and cucurbit beetles residing in the humid wilted floral chambers. In conclusion, our findings showcase squash flowers as a humid microhabitat that influences reproductive success directly by affecting pollen viability and indirectly by altering interactions with squash bees, their specialist pollinators.

## Introduction

The stunning evolutionary diversity of floral display has long been attributed to the sensory and cognitive biases of animal pollinators as agents of natural selection ^1–5^. However, flowers are more than “sensory billboards”, advertising nectar, pollen and other rewards to pollinators through conspicuous floral signals ^6^. Additional selective forces, comprising biotic (herbivores, florivores, pathogens) and abiotic (wind, rain, solar radiation) factors, contribute to the evolution of floral form and function ^7–9^. Many floral traits are now understood to play defensive roles against antagonists, including rhythmic floral closure ^10^, sticky exudates or trichomes ^11^, herbivore-repellent or antifungal pigments ^12^, toxic nectar ^13^ and organ-specific antibacterial volatiles ^14^. Abiotic factors also can shape components of floral traits, from bracts and pigments that protect pollen from rain ^15^ or UV exposure ^16^ to solar tracking ^17,18^ or thermogenic properties ^19^ that provide warmer environments for pollinators and growing pollen tubes ^20^. Flowers also host diverse ‘anthophiles’ (flower-associated organisms), including mutualists, commensals, predators, and microbiomes ^21–23^. In these examples, flowers function as miniature nurseries containing multi-trophic communities, highlighting the importance of flowers as microcosms. Despite growing evidence linking floral microclimates to reproductive success, few studies test explicit hypotheses regarding both direct and indirect impacts of floral microclimate on plant fitness.

The floral microclimate is defined by the measurable physical conditions from the flower’s surface across its enclosed headspace (micrometeorology) ^24–27^. Developing flower buds contain air spaces that are insulated from the external environment by the perianth. Upon floral opening, this microclimate is impacted by floral shape, orientation, pigmentation, trichomes or cuticular waxes ^28–31^. At present, floral temperature is the best-studied aspect of floral microclimate, especially in thermogenic flowers that generate endogenous heat well above ambient temperatures ^32–34^. Thermogenicity has evolved in at least 12 angiosperm families and two cycad families, potentially existing in Cycadales for at least 200 million years ^35^. Thermogenic plants are often associated with strongly scented floral chambers or cycad cones in which pollinators obtain food, mates, brood sites or shelter ^36–40^. More common, however, are flowers that passively absorb heat via parabolic shapes, inflated calyces, dark pigments, or sun tracking ^28,30,31,41^. Regardless of the mechanism, warmer flowers benefit both plants and pollinators ^28^. Less widely appreciated is that many flowers maintain more humid headspace air than their surroundings due to nectar evaporation, leaky petal cuticles ^42^, or transpiration ^43–45^.

Corbet et al.^46^ measured floral humidity and its influence on nectar equilibration rates, suggesting potential impacts on pollinator preferences. Subsequently, von Arx et al.^44^ showed that ephemeral humidity gradients from nectar evaporation and transpiration in *Oenothera cespitosa* flowers can serve as nectar-presence cues for their hawkmoth pollinator *Hyles lineata*. In *Datura wrightii*, headspace humidity gradients are ten times greater than those reported in other species ^45^ and originate primarily from corolla transpiration rather than nectar evaporation, making them persistent. Similar to *H. lineata*, the nocturnal hawkmoth *Manduca sexta* prefers humidity-enhanced artificial flowers over ambient conditions and can use floral humidity for efficient nectar extraction^45^. While these studies establish that flowers with large petal surface areas and copious nectar volumes can produce efficacious humidity gradients, their broader ecological functions beyond pollinator attraction remain poorly understood in both wild and domesticated plants.

Here, we explored the floral microclimate and its contributions to plant-pollinator interactions in the blossoms of squash and pumpkin plants (*Cucurbita* spp; Cucurbitaceae). The domestication and expansion of squash from its ancestral range in Mesoamerica into North America, along with its mutualistic relationship with squash bee pollinators, make cucurbits a valuable system for studying traits influenced by natural and artificial selection ^47–49^. In addition to the global economic importance of squash-related crops ^50^, squash flowers provide rich nectar and pollen resources to both specialist and generalist bee pollinators. Squash plants are monoecious, with separate staminate and pistillate flowers on the same plant; pistillate flowers provide no pollen but accumulate three times more nectar than staminate flowers in *C. pepo* ^51^. Thus, bee visitation is required to transfer pollen between flowers and achieve fertilization.

Specialist squash bees, *Xenoglossa (*formerly *Peponapis* or *Eucera* ^52^*) pruinosa* (Apidae), engage in a mutualistic relationship with squash flowers. Female squash bees provision larvae with pollen, while both sexes consume floral nectar ^53^. Importantly, while female *X. pruinosa* bees are ground-nesting, male bees use *Cucurbita pepo* flowers as shelters, mating sites, and food sources ^54–59^. In contrast, generalist bees, including female bumblebees (*Bombus impatiens*) and honeybees (*Apis mellifera*), visit squash for nectar but do not reside in the flowers. Generalist bees are important for squash pollination, especially beyond the native range of squash bees ^51 b,60,61,62 a^. However, bees are not the only visitors to squash blossoms. Specialist herbivores, such as spotted (*Acalymma vittatum*) and striped cucurbit beetles (*Diabrotica undecimpunctata*; Chrysomelidae), feed on squash and gourd flowers ^63–66^. Their relationship with squash is antagonistic, as the beetles feed, mate, and defecate within the flower, often in large numbers, consuming floral tissues and spreading harmful *Erwinia* bacterial pathogens ^67^. The fact that both mutualists (male squash bees) and antagonists (cucurbit beetles) frequently reside in squash flowers for extended periods suggests that the floral microclimate may play both direct and indirect roles in squash reproductive fitness. However, it remains unclear whether floral temperature or humidity affects the visitation and habitat of squash flowers.

We assessed the influence of floral temperature (T) and relative humidity (RH) on pollen survival and insect visitation in both wild and domesticated squash flowers. First, we monitored floral T, RH, and insect visitors in a population of wild buffalo gourd (*Cucurbita foetidissima*) in its natural habitat. Next, we established baseline data for floral T and RH in cultivated squash (*Cucurbita pepo*) across agricultural fields, with a focus on how floral T is affected by passive solar radiation during floral senescence, when the corollas wilt and form closed chambers. Third, we experimentally examined the sources of floral RH gradients and their effects on pollen tube growth and bursting using greenhouse-grown squash (*C. pepo* subsp. *pepo*). Finally, we conducted field experiments, to evaluate how florivory by beetles impacts floral RH, insect visitation, and residence times within squash floral chambers.

## Results

Wild and domesticated squash flowers show similar micrometeorological features The flowers of wild squash *Cucurbita foetidissima* are large, with a floral volume of 31.16 ± 4.94 cm^3^ for males (mean ± SD, n=6), and 38.42 ± 17.11 cm^3^ for females (mean ± SD, n=7). They attract a variety of insects for nectar or pollen, while others take residence in the floral chamber (Fig. 1A). Only two of the 23 flowers observed along a transect within a natural population of *C. foetidissima* near Saguarita, Arizona, USA (see Methods for details) were unoccupied. Most flowers were inhabited by aggregations of *Drosophila pseudoobscura* flies (Video S1). Additionally, six of the 21 occupied flowers hosted chrysomelid cucumber beetles, three flowers were inhabited by *Xenoglossa pruinosa* squash bees and two flowers were visited by *Xenoglossa strenua* squash bees.

**Fig. 1.**
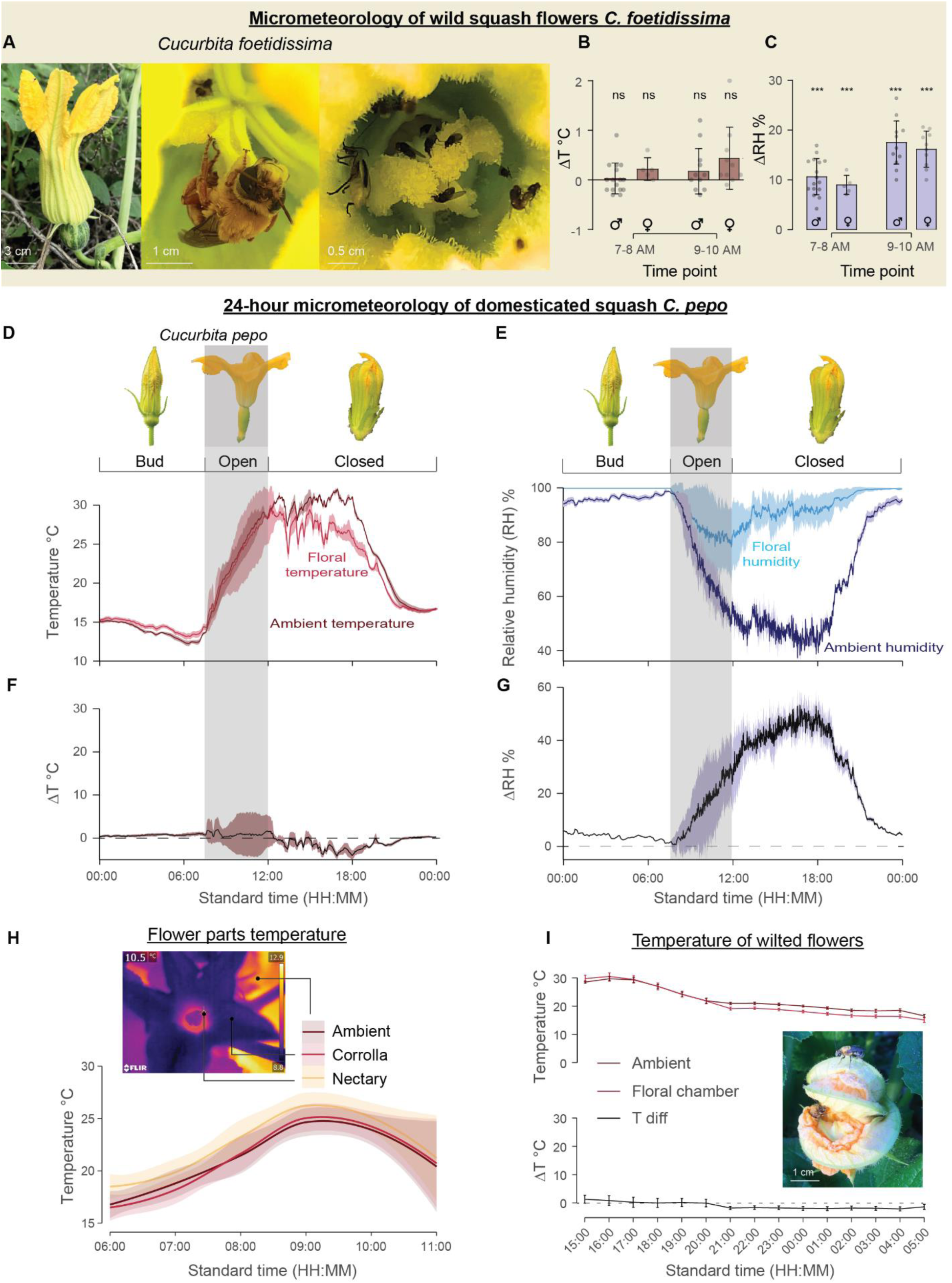
Wild and domesticated squash flowers maintain a humid microclimate within their headspace. A. Images of wild squash *C. foetidissima* depict several insects inhabiting the floral chamber. Scale bar at the bottom left corner. B. Micrometeorology of wild squash *C. foetidissima*, highlighting the difference in temperature between the floral chamber and the ambient environment (ΔT) at two distinct time points during the morning hours. Dot plots display individual flowers overlaid with histograms showing the mean ± SD. “ns” indicates not significant, and “***” denotes P<0.0001 (One-sample t-test against zero). Icons indicate flower sex C. Same as b but highlighting the difference in humidity between the floral chamber and the ambient (ΔRH). D. 24-hour measurements of the floral and ambient temperature across three stages of *C. pepo* flower development, as depicted in the images on top. The gray bar highlights a brief window of approximately 6 hours when the flowers stay open. Solid lines represent the mean, with the shaded area indicating the SD around the mean of n=4 flowers. E. Same as D but showing floral humidity across 24 hours. F. The ΔT *C. pepo* flowers extracted from the difference between ambient and floral T shown in panel D. G. The ΔRH *C. pepo* flowers extracted from the difference between ambient and floral RH shown in panel E H. Temperature of flower organs measured hourly during the open phase from 6:00 to 11:00 using a FLIR thermal camera. A representative image with pointers at different flower locations is displayed on top. Solid lines show smooth splines going through means for every time point (n=10). The shaded area represents the 95% confidence interval around the mean. I. Floral and ambient temperature during the closed phase of the flower (top) and the ΔT (bottom) for n=21-35 flowers. Color-coded lines represent the mean and the vertical bars show SE for each time point. The dashed line in the bottom panel highlights no difference in floral and ambient temperature. The inset image shows male squash bees exiting a wilted flower in the morning after sleeping overnight in the floral chamber.

The temperature (T) of both male and female flowers did not differ from ambient T (Fig. 1B), but floral relative humidity (RH) was notably higher than ambient levels (Fig. 1C). Despite high ambient RH of 79.30 ± 3.33 %RH (mean ± SD; VPD= 0.41 kPa) during the first time point at 7-8 am, the floral RH exceeded ambient by 10.25±3.34% RH (Fig. 1C; male flowers: t=11.70, df=15, P<0.0001; female flowers: t=10.64, df= 4, P=0.0004). In contrast, floral T was nearly identical to ambient T, with only a 0.1°C difference between them (ambient temperature= 22.8 ± 0.62°C; male: t=0.40, df= 15, P=0.69; female: t=2.15, df=4, P=0.09). By the second time point between 9-10 am, ambient humidity dropped to 64.58 ± 6.78 %RH (VPD=1.41 kPa), but the floral ΔRH was 16.94 ± 4.02% (Fig. 1C; male: t=15.11, df=13, P<0.0001; female: t=14.12, df=9, P<0.0001). Although ambient T quickly increased to 28.95 ± 1.85°C by 10:00 hrs, the floral T did not differ significantly from ambient levels (Fig. 1B; male: t=1.53, df=15, P=0.14; female: t=2.22, df=4, P=0.053) suggesting that wild squash flowers regulate headspace RH but not T during the open phase of the flower.

The flowers of *Cucurbita pepo,* a domesticated variety of squash, are comparably large, with male and female flower volumes of 29.30 ± 7.34 cm^3^ and 41.90 ± 19.83 cm^3^, respectively (Table S8). We characterized the floral microclimate of *C. pepo* over an extended period with greater temporal resolution, utilizing a squash farm established near University of Guelph, Ontario, Canada. The floral temperatures of *C. pepo* closely tracked ambient levels (Fig. 1D). Despite a substantial (∼15°C) difference between morning and late afternoon ambient T, the difference between ambient and floral temperature (ΔT) was nearly zero (Fig. 1F). Conversely, floral RH consistently remained higher than background levels, regardless of ambient humidity (Fig. 1E). In the bud stage, when the flower is tightly closed, the headspace humidity is saturated at 100% RH. As the flower opens at sunrise, floral RH dissipates to the much lower ambient RH (Fig. 1E). Once the flower closes around noon, floral RH within the chamber increases and the ΔRH peaks around 18:00 hrs when ambient RH drops to its lowest point (Fig. 1G). In summary, both wild and domesticated squash plants maintain a humid microhabitat in their large volumes of floral headspace, without comparably large differences in floral temperature.

FLIR thermal camera images (Fig. 1H) revealed that flowers of both sexes showed identical T patterns (LMM: t=-0.85, df=409.02, P=0.39) during the open phase. Corolla and ambient T also were identical (t= -0.13, df=408.99, P=0.89). The nectary was slightly warmer than the other two zones across all time points but not significantly warmer than the ambient or the corolla T (t=1.35, df= 408.99, P=0.17). For wilted and closed flowers, the T were similar to ambient air T. In general, there was a slight trend for the flowers to be warmer than ambient during the day but cooler than ambient at night (Fig. 1I). However, the ΔT was close to 0° C, suggesting that difference is likely a result of passive heating and cooling of flowers and that wilted flowers are not warmer during the night, as is observed for thermogenic flowers ^84,85^. Altogether, these results show that squash flowers do not regulate floral T but maintain humidity for the entirety of the flower’s life.

Floral transpiration, but not nectar, is the major source of floral humidity We evaluated the relative contributions of nectar evaporation and cuticular surface to floral humidity gradients along a vertical transect beginning near the floral nectary.

Squash flowers present a steep increasing gradient of humidity from the ambient air outside the flower to the nectary at the flower base (Fig 2A). These data were qualitatively consistent for both male and female flowers of *C. pepo* (Fig 2A and B). At the flower base, intact control female flowers were 34.71 ± 2.03% RH (mean ±SE) above ambient humidity (henceforth ΔRH), and males were 32.91 ± 3.09% ΔRH. At the threshold of the flower (approx. 60 mm on the transect) floral ΔRH is 4.66 ± 0.72% for female flowers and 4.23 ± 1.67 % for male flowers. At the end of the vertical transect, approximately 140 mm away from the nectary, floral ΔRH is nearly ambient with 2.96 ± 0.47 % and 3.27 ± 0.55 % for female and male flowers, respectively.

**Fig. 2:**
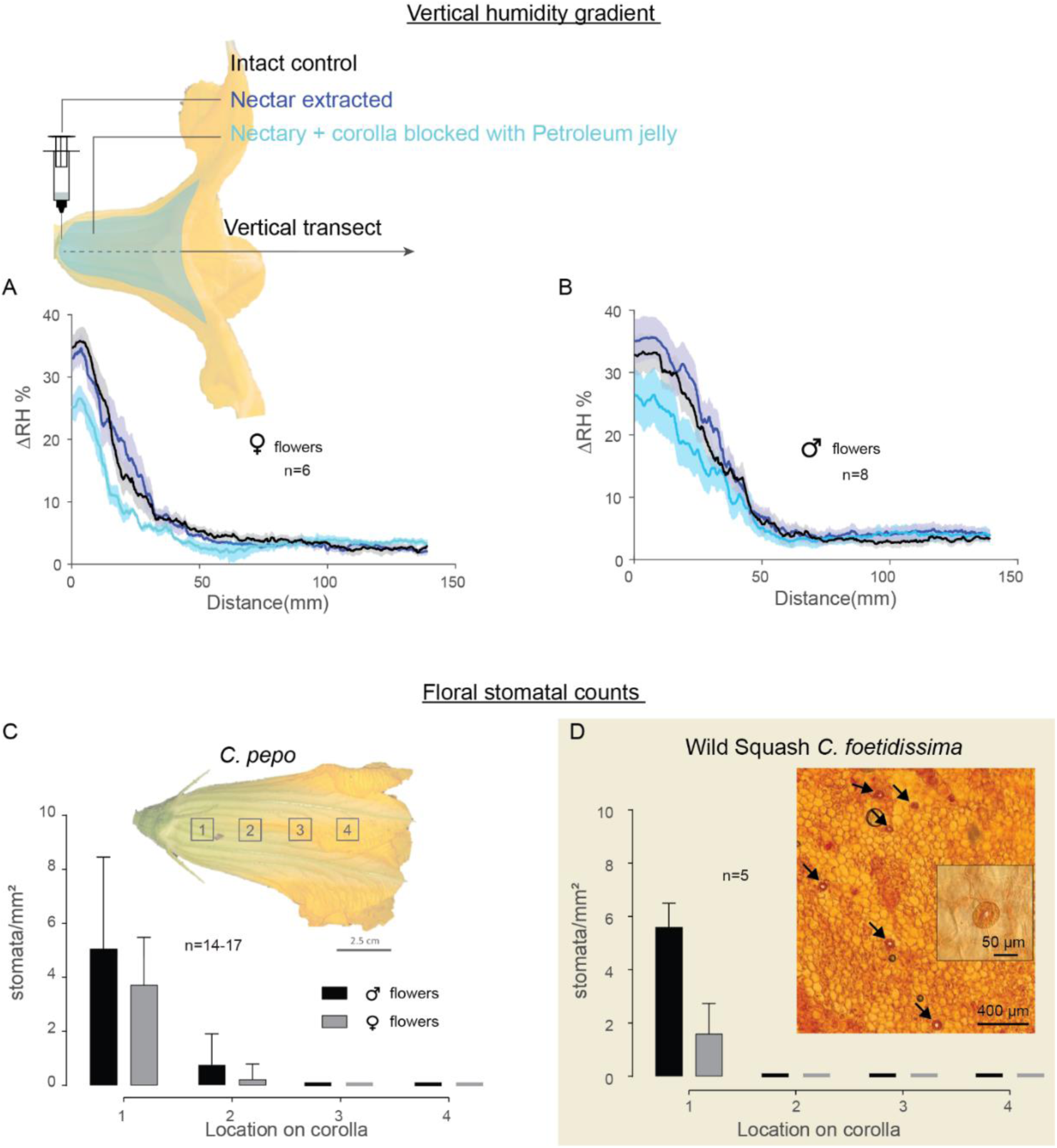
Floral stomata and nectar contribute to the floral humidity gradient within squash headspace. A. Vertical transect depict floral humidity in male *C. pepo* for unmanipulated controls, nectary blocked, and nectary + corolla blocked conditions. The top schematic illustrates a transect for measuring the vertical humidity gradient, with the syringe indicating nectar extraction and the cyan-shaded area representing Petroleum jelly application to coat the adaxial surface. Colors denote different treatments, and sample sizes are provided within the panel. Solid lines and shaded areas represent the mean ± SE. Vertical gradients are compared statistically using a nonlinear exponential decay model. Comparisons of model parameters are presented in Table S1 and S2. B. Same as A, but for female flowers. C. and D, Stomata on the inner surface of *C. pepo* and *C. foetidissima* (Also see Fig. S1C and S1D). Bar graphs depict the mean ± SD of stomata number/mm^2^ sampled at four locations on the inner surface of the flower as indicated in the inset image in panel c. Black bars for male and gray for female flowers. The inset image in panel d shows a Safranin-stained floral peel pictured under a microscope at 10x. Arrows indicate stomata observed at location 4 of the petal. An enlarged image of a floral stoma with visible chloroplasts in the guard cells is included. The species identity is shown at the top of the panel, scale bars are provided in the bottom right corner of the image, and sample sizes are mentioned in each panel.

Experimental nectar removal influenced the floral RH gradient. Multiple comparisons across treatments for the two model parameters α for the decay rate and y0 for intercept suggested that both parameters differed significantly between the intact control and nectar-extracted male and female flowers (Table S1 and S2). Furthermore, flowers with their stomates and nectaries blocked by the application of petroleum jelly showed a substantial reduction in the difference of humidity (ΔRH 25.6 ± 2.51 %, mean ±SE ;) from ambient levels compared with the other two treatments (Fig. 2A & 2B and Table S1 and S2). The decay rate α of floral RH was steeper for petroleum jelly-smeared flowers than for nectar-extracted (female flowers: *t*= -15.56, *P*<.0001; male flowers: t= 11.52, P<0.0001) and intact control flowers (female flowers: *t*= -9.58, *P*<0.0001; male flowers: t= 27.39, P<0.0001). The intercept y0 for the petroleum jelly-treated flowers was significantly lower than the intercepts of nectar-extracted (female flowers: *t*= 21.08, *P*<0.0001; male flowers: t= 43.79, P<0.0001) and intact control flowers (female flowers: *t*= 26.99, *P*<0.0001; male flowers: t= 33.41, P<0.0001).

Domestic and wild squash flowers show the same pattern of stomatal distribution Peels from locations 1 and 2 of the adaxial surface of the corolla revealed the presence of multicellular trichomes within the floral chamber (Fig. S1A). Additionally, observations from peels indicated that flowers receive water and nutrients through the xylem and protoxylem (Fig. S1B). Stomata were abundant on the surfaces of both *C. pepo* and *C. foetidissima* flowers (Fig. 2C & 2D). The distribution of stomata matched the vertical floral humidity gradient, with the highest density corresponding to the highest humidity location near the nectary. Stomatal density decreased markedly towards the corolla limb (locations 3 and 4), where stomata were essentially absent. Male flowers typically exhibited a higher stomatal density than did female flowers. This distribution pattern of stomata aligns with observations made in other species of cultivated squashes (Fig. S1C and S1D for *C. maxima* and *C. moschata*) and together these findings suggest that squash floral humidity is not solely a passive consequence of nectar evaporation.

Reduced floral humidity diminishes pollen viability.

We used a common garden greenhouse planted with *C. pepo* to test whether experimental reduction of floral humidity in male squash flowers negatively impacts pollen viability (Fig. 3A).

**Fig. 3.**
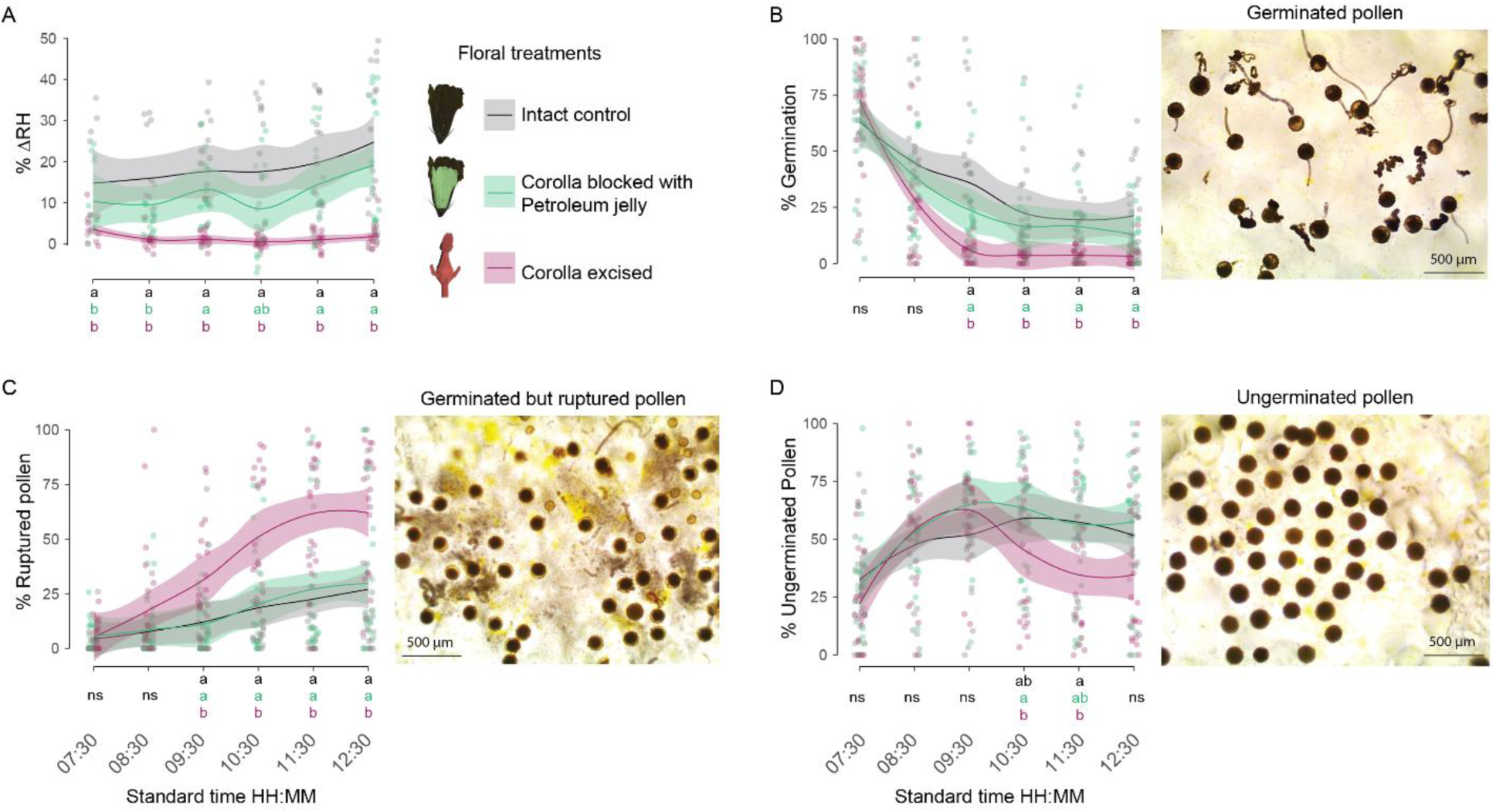
Reduced floral humidity diminishes pollen viability and increases the incidence of pollen bursting in greenhouse squash flowers. A. Hourly measurements of floral humidity in squash flowers were taken for intact control flowers, flowers treated with Petroleum jelly on the inner surface, and flowers with corolla excised. B. Percentage of pollen germinated on Brewbaker Kwack ^69^ medium for flowers subjected to the two treatments and intact control conditions. C. Percentage of pollen rupturing on the germination medium. D. Percentage of pollen remaining ungerminated on the medium. Example images are provided on the right of each panel. Treatments are color-coded, and legends depict icons for each treatment. The solid lines represent the mean plotted using LOESS method, with the shaded area indicating a 95% confidence interval around the mean. Statistical significance is indicated at the bottom of each panel, color-coded by treatments. Common letters denote P>0.05, while distinct letters signify P<0.05, and ‘ns’ for not significant ANOVAs. Refer to Table S3 for pairwise comparisons for each time point.

The floral ΔRH differed significantly between treatments (ANOVA, F=68.03, df=2, P<0.0001) and across time (F=3.38, df=5, P=0.005). Flowers with their adaxial surfaces coated with petroleum jelly exhibited consistently lower ΔRH than intact control flowers (P=0.0001), albeit significantly higher than flowers with excised corollas (P<0.0001) for most time points (Fig. 3A and Table S3). As expected, corolla-excised flowers displayed humidity levels nearly equivalent to ambient conditions (see Fig. 3A), significantly lower ΔRH than intact control flowers for all time points (P<0.05) and Petroleum jelly-coated flowers for time points at 09:30, 11:30, and 12:30 hrs (Table S3).

The observed decreases in floral RH were associated with significant differences in pollen germination rates between treatments (ANOVA: F=27.11, P<0.0001) and across time (F=53.22, P<0.0001). Overall, pollen germinability decreased over time, even within intact control flowers (Fig. 3B). At anthesis, 65-70% of the pollen was viable in all three treatments. However, by floral closure at midday, 20% of the pollen remained viable in control flowers, 12% in petroleum jelly-treated flowers, and only 3% in corolla-excised flowers (Fig. 3B). Multiple comparisons between treatments indicated significantly lower germination rates in pollen from the corolla-excised treatment compared with control and petroleum jelly-treated flowers for all time points after 08:30 hrs (P<0.01, Table S3). There was insufficient evidence for a difference in pollen germination rates between control and petroleum jelly-treated flowers (P>0.05, Table S3).

Additionally, a portion of the pollen germinated but subsequently developed ruptured pollen tubes, leaving substantial debris in the germination medium (Fig. 3C). There was a significant difference in the proportion of ruptured pollen between treatments (F=30.80, df=2, P<0.0001), with the proportion increasing over time (F=23.11, df=5, P<0.0001). Post hoc tests revealed a notably higher bursting rate in pollen from corolla-excised flowers compared with control (P<0.01) and petroleum jelly-coated flowers (P<0.01). Specifically, the bursting rate increased significantly after 08:30 hrs for pollen in the corolla-excised treatment (Supplementary Table S3). There was no compelling evidence for a difference in pollen bursting rate between control and petroleum jelly-treated flowers at any time point (ANOVA: P=0.78, Table S3).

Moreover, a portion of the pollen would not germinate, with inviable proportions differing significantly between treatments (F=7.23, df=2, P=0.0008) and overtime (F=11.64, df=5, P<0.0001). The viable pollen proportion was significantly higher in control and Petroleum jelly-treated flowers than in the corolla-excised treatment across time (P=0.009 & P=0.001, respectively). Specifically, the time points at 10:30 and 11:30 hrs differed significantly between treatments (Fig. 3D & Table S3). The difference between control and petroleum jelly-coated flowers was negligible (ANOVA: P=0.81, Table S3) throughout the experiment.

Floral micrometeorology affects the visitation of specialist pollinators of squash flowers but not the generalist pollinators

To simulate beetle-induced damage under controlled conditions, we introduced perforations either in the floral chamber or on the flower limb and compared their impacts on humidity and temperature measurements. Our findings revealed that temperature variation across floral treatments and period was insignificant (Fig. 4D, Table S4), consistent with (Fig. 1). However, floral ΔRH varied significantly between treatments (Table S4) and showed a general trend of increase as the ambient conditions became warmer and drier during the latter part of the morning (Periods 2 & 3; Table S4). Flowers with chamber damage exhibited markedly lower humidity levels than those with limb damage and the undamaged control flowers across all periods (LMM: P<0.0001; Fig. 4C). Corolla limb damage did not alter humidity levels compared with the undamaged control flowers for either period (LMM: P=0.4860; Fig. 4C).

**Fig. 4.**
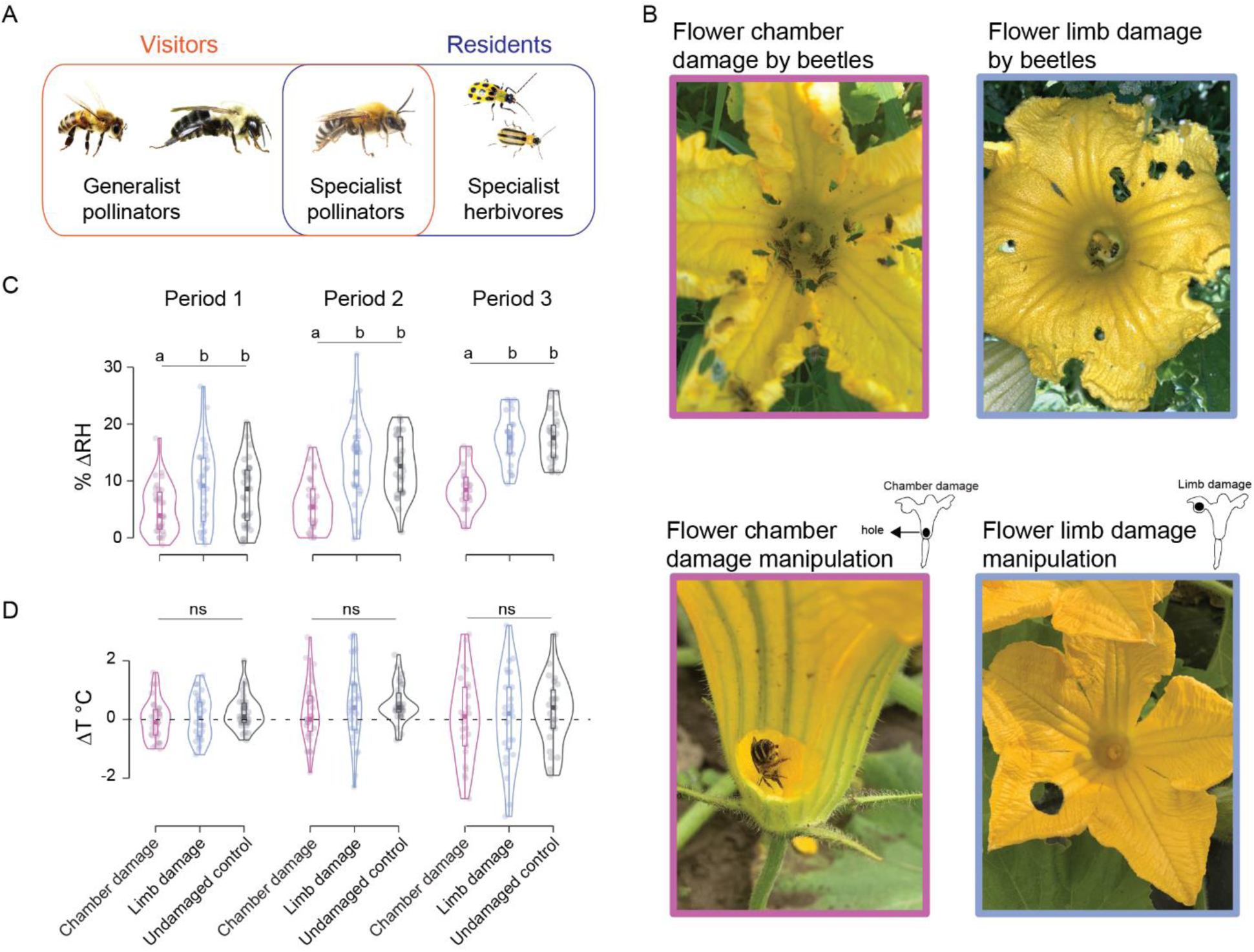
Floral chamber damage impacts humidity but not temperature in squash flowers. A. Illustrating the visitors and residents of squash flowers. Bumblebees and honeybees serve as generalist pollinators, while squash bees specialize in pollinating squash varieties. All bees are regarded as mutualists in this ecosystem due to their pollination contribution whereas Cucurbit beetles are considered antagonists here ^64^. B. Images depict two patterns of beetle damage (Top row). Damage to the floral chamber is likely to reduce floral humidity through leakage, while damage to the corolla limb may not have the same effect. Colors surrounding the images serve as guides for experimental manipulations depicted in the bottom row, replicating the impact of floral humidity on visitors and residents of squash. Icons at the bottom represent the treatments and are used in subsequent figures instead of the images. C. Violin plots with embedded boxplots display the floral ΔRH (relative humidity) and ΔT of flowers under two experimentally manipulated treatments, along with undamaged control flowers, across three observation periods per day. Period 1 corresponds to the early morning hours (7-8:30 am); Period 2 to (9-10 am); Period 3 to (10-11 am). Colors correspond to different treatments, and letters above the violin plots indicate significant differences at P<0.05 (LMM, Table S4). A dashed line at ΔT 0°C indicates no difference from ambient T.

Leveraging these differences in floral RH across treatments, we tested whether they affect the behavior of both mutualists and antagonists towards domestic squash flowers. In the first experiment, we focused on evaluating the impact on bees as floral visitors. We observed a decreased overall visitation frequency of bees on chamber-damaged flowers compared with both limb-damaged (GLMM: z= 3.11, P=0.001) and undamaged control flowers (GLMM: z=3.03, P=0.002; see Fig. S2A for a summary of all bee species). Bee visitation increased progressively from period 1 to period 3 (GLMM: z=6.35, P<0.001), with no significant interaction effect between observation period and treatment (P>0.05). A post hoc analysis comparing the means of each treatment within individual periods revealed a significant impact of bee visitation in periods 1 and 2, but not in period 3. Despite the effect of floral humidity manipulation on bee visitation frequency, there were no discernible differences in the foraging duration once bees alighted on the flowers (Fig. S2B and Table S5).

Subsequently, we scrutinized the data according to bee species to ascertain whether the observed differences were attributable to specific bee species or were representative across all visiting bees. In general, squash bees exhibited lower visitation frequencies on chamber-damaged flowers compared with limb-damaged (Fig. 5B; GLMM: z= 3.35, P=0.0008) and control flowers (GLMM: z= 3.04, P=0.002). We observed a significant effect of observation periods 2 and 3 compared with period 1, along with some indication of an interaction effect between limb-damaged and undamaged flowers with period 3 (Table S5). However, there was no evidence of an effect on the foraging duration of squash bees across any of the flower treatments (Fig. 5C and Table S6).

**Fig. 5.**
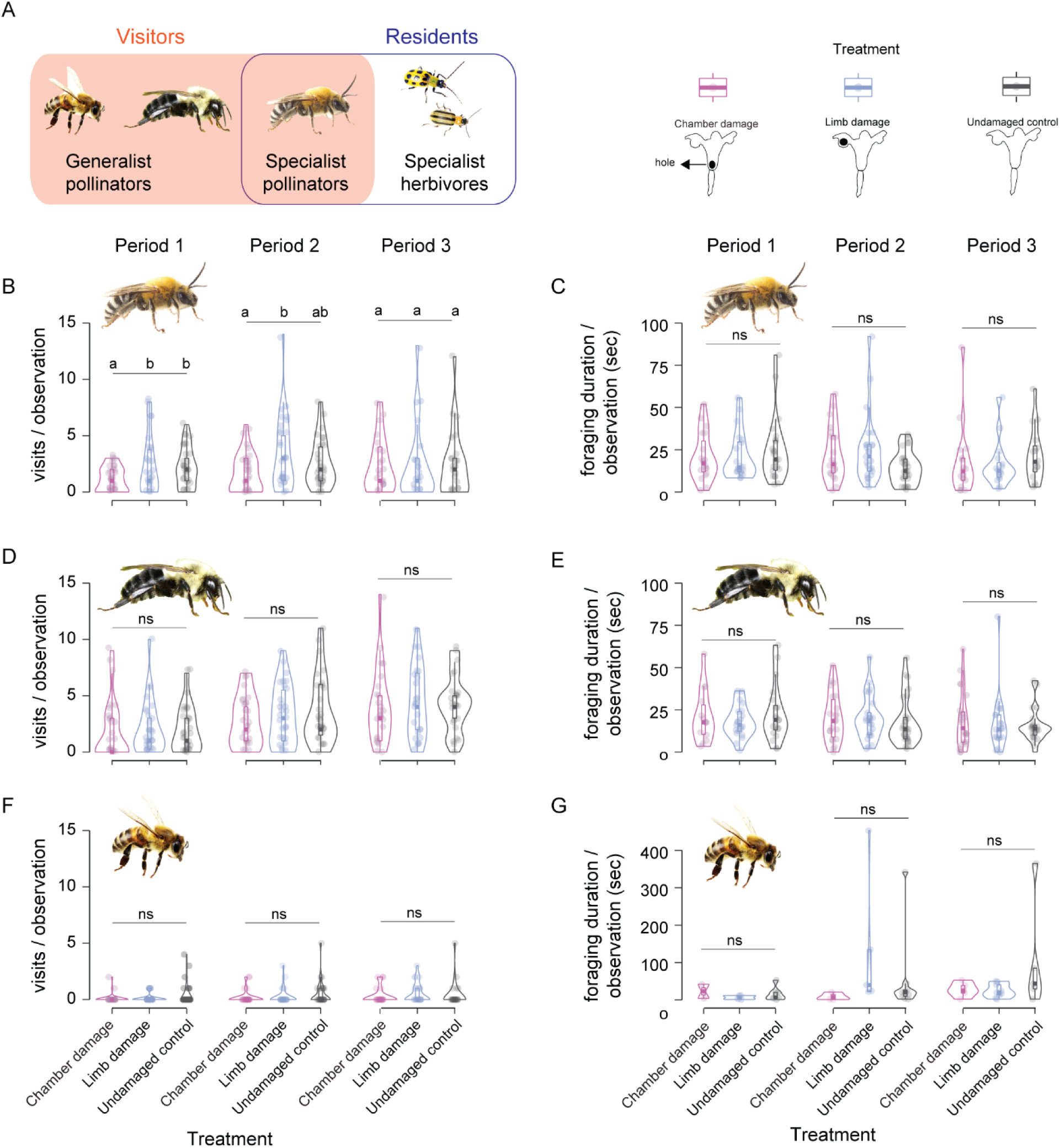
Floral humidity largely affects visitation by specialist pollinators. A. The highlighted portion in the summary figure indicates the visitor data shown here, reflecting generalist and specialist pollinators in the foraging context. B. D, F, Violin plots with embedded boxplots illustrating the visitation frequency of Squash bees, bumblebees, and honeybees per 10 minutes of observation across Periods 1-3 to flowers with different treatments (refer to icons at the top left). C. E, G, Violin plots with embedded boxplots demonstrating the foraging times of Squash bees, bumblebees, and honeybees per 10 minutes of observation across Periods 1-3 to flowers with different treatments (refer to icons at the top left). All treatments are color-coded. Different letters indicate groups that are significantly different within a period (GLM, P<0.05); “ns” indicates not significant.

There were no pronounced differences in bumblebee visitation associated with any flower treatment (Fig. 5D). Comparisons between limb-damaged and chamber-damaged flowers (GLMM: z=0.93, P=0.34) and between chamber-damaged and undamaged control flowers (GLMM: z=0.55, P=0.57) were both statistically insignificant. Bumblebee visitation frequency doubled from Periods 1 to 3, contrasting with the relatively uniform visitation pattern of squash bees across all periods (Fig. 5D). There was no discernible effect of floral treatments on bee foraging duration across observation periods (Fig. 5E, Table S6). However, bumblebee foraging duration significantly increased from Period 1 to 2 (Table S6, P=0.01), with a significant interaction effect of undamaged flowers observed in Period 2 (Table S6). Overall, bumblebee foraging duration slightly decreased from Periods 1 through 3, while their visitation frequency increased with time (Fig. 5D).

Honeybees were less common than bumblebees and squash bees as flower visitors, with no discernible differences in their visitation frequency across floral treatments (Fig. 5F, Table S5). Similarly, there was no disparity in their foraging duration on either type of flower (Fig. 5G, Table S6). These findings indicate that flowers with reduced humidity are likely to attract fewer visits from the specialist pollinator *X. pruinosa*.

In the second experiment, we tested whether floral manipulations and their impacts on RH affect specialized insects that use faded, closed flowers as shelters. Specialist cucurbit beetles were observed in 81.3% of the investigated flowers (Fig. 6B), and flowers often housed tens of beetles together (Video S2). Among the occupied flowers, the relative percentages of occupancy were as follows: chamber-damaged flowers (30.1%) < limb-damaged flowers (33.2%) < undamaged control flowers (36.7%; Fig. 6B). Overall, more beetles were found inhabiting undamaged control flowers than the chamber-damaged (emmeans: P<0.0001) or limb-damaged flowers (emmeans: P=0.04; Fig. 6C). However, beetle occurrence was similar between chamber-damaged and limb-damaged flowers (emmeans: P=0.06). While farm sites did not affect beetle residence in flowers (GLM, z=-1.14, P=0.25), the squash variety exerted a strong influence, with flowers of *C. maxima* attracting significantly more beetles than flowers of *C. moschata* and *C. pepo* (Fig. S3 and Table S7).

**Fig. 6.**
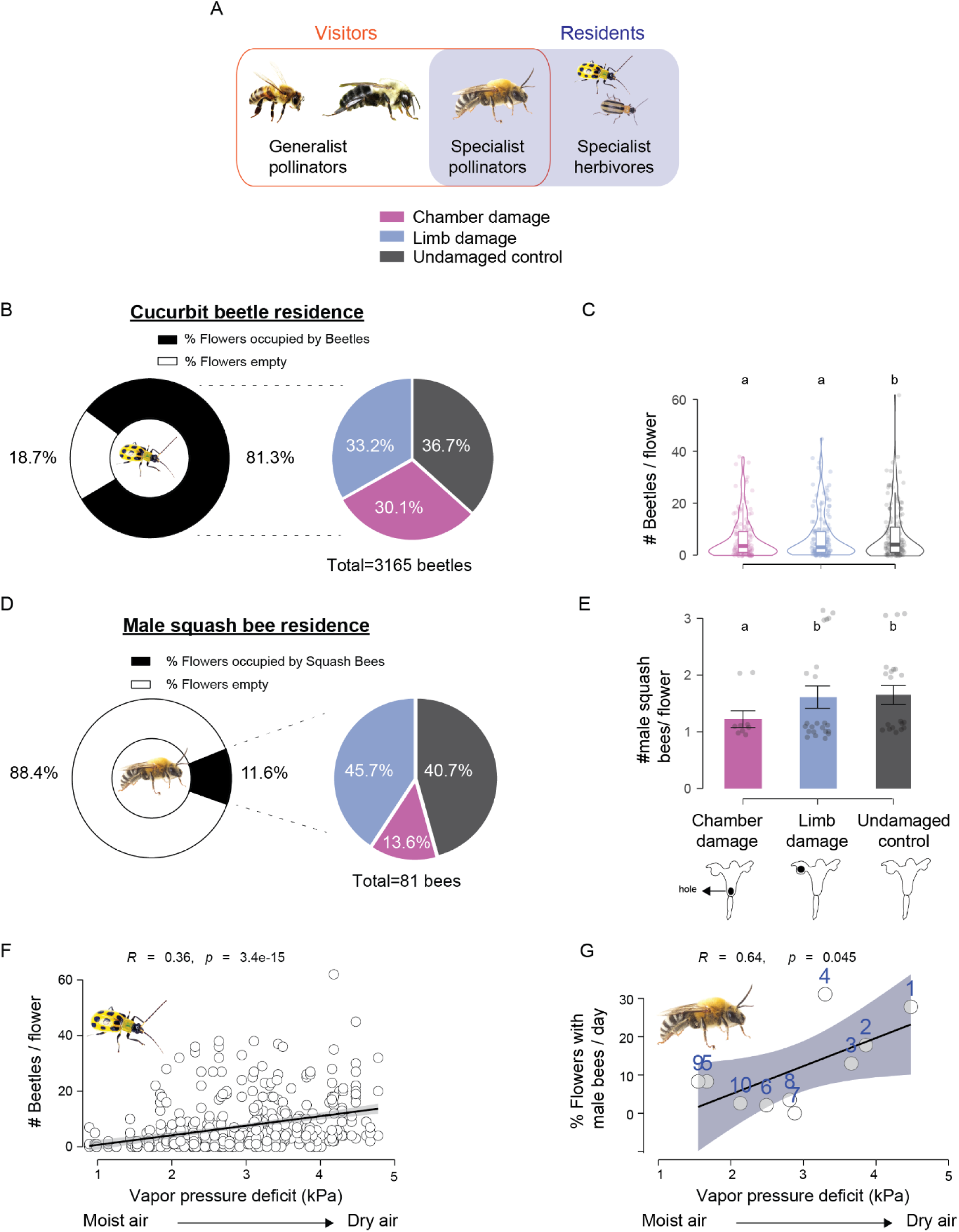
Floral humidity manipulations affect squash bee residence in wilted flowers. A. The highlighted portion in the summary figure indicates the resident data shown here, reflecting squash bees and cucurbit beetles. B. Pie charts presenting Cucurbit beetle residence data. The pie chart on the left shows overall beetle occurrence across the 10 days of the sampling period. The chart on the right shows the proportion of beetles within chamber-damaged, limb-damaged, and undamaged-control flowers. C. Violin plots with embedded boxplots displaying the number of beetles per flower across the floral treatments. Letters on top of the panel indicate significant differences (GLM, P>0.05). D. Pie charts depicting male squash bee residence data in the same format as panel b. E. Number of male squash bees per flower (Data included for only the flowers with bees because they are rare. Often more than one bee inhabits a single flower). F. f and g) A correlation between the number of beetles per flower plotted against the vapor pressure deficit of the ambient air (VPD) in (f) and the percentage sampled flowers occupied by male squash bees plotted against the VPD in (g). The correlation coefficients and the P values are shown in each panel. Shaded area around the regression line indicates 95% confidence interval. The numbers on the datapoints in panel g reflect the sampling date, with 1 being the first in the season, and 10 being the last sampling date. Shaded area indicates 95% confidence interval.

Male squash bees were frequently observed sharing flowers with cucurbit beetles, and often a single flower accommodated more than one bee. However, only 11.6% of the sampled flowers contained male squash bees (Fig. 6d). Within the flowers hosting male bees, the relative percentages of occupancy were as follows: chamber-damaged (13.6%) < limb-damaged (45.7%) < undamaged control (40.7%; Fig. 6D). Overall, chamber-damaged flowers exhibited significantly lower male squash bee residence compared with limb-damaged (Fig. 6E; zero-inflated GLM, z=2.89, P=0.0038) and undamaged control flowers (zero-inflated GLM, z= 2.51, P=0.01). Squash variety did not impact male squash bee residence (zero-inflated GLM, z= -1.29, P=0.19).

Finally, we explored how the occurrence of residents in flowers correlates with the dryness of the air (Vapor pressure deficit). A correlation between the number of beetles in flowers and the vapor pressure deficit revealed that beetles more frequently inhabit flowers when the ambient air is drier (Fig. 6F; t=8.16, df=448, P<0.0001). Regarding squash bees, since their presence in flowers usually ranges from one to two individuals, we calculated the percentage of flowers occupied by bees for each observation day rather than the absolute number of bees in each flower. A higher percentage of flowers was occupied by bees when the ambient air was drier (Fig. 6G; t=2.37, df=8, P=0.045).

## Discussion

Plant micrometeorology is an emerging field of science that considers the interactions of plant organs with their surrounding environment^25,29^. Micrometeorological studies on flowers have mostly focused on thermogenic or thermoregulating flowers like water lilies, lotus, and skunk cabbage that create warm microhabitats favored by their pollinators over cooler ambient conditions ^84–87^. Despite their ecological significance, thermogenic flowers are rare, occurring in only 12 of 416 angiosperm families ^35^. In contrast, above-ambient humidity is much more widespread in flowers ^43,88^, but its importance in providing microhabitats for flower-inhabiting organisms (anthophiles) remains understudied ^23^. We examined the microclimate within the large blossoms of both domesticated and wild squash species, which are frequented by and serve as shelters for many insects. Our findings indicate that squash flowers maintain significantly higher relative humidity (RH) levels than ambient conditions throughout a 24-hour period, while floral temperatures (T) remain consistent with ambient T (Fig. 1). Our data suggest direct impacts of floral RH on plant fitness, as evidenced by a higher incidence of pollen rupture in humidity-leaked flowers relative to control flowers with typical RH levels (Fig. 3). Field assays demonstrated that altering floral RH influences interactions with both mutualistic squash bees and antagonistic cucurbit beetles. RH manipulation reduced the landing frequency of squash bees during foraging but did not alter the duration of their foraging once landed (Fig. 5). Additionally, wilted flowers with manipulated RH were less frequently inhabited by squash bees compared with sham controls and unmanipulated flowers (Fig. 6). Overall, these findings suggest that the RH of squash flowers influences the specialist insects in this system, portraying squash flowers as a humid microhabitat for these small insects.

### Micrometeorology of Squash flowers

Thermogenic flowers often feature large internal chambers that retain heat ^89^, a trait shared by some flowers with high RH levels, such as *Datura wrightii*, which are large (8.44 ± 0.58 g fresh weight) and voluminous (83.95 ± 14.40 cm^3^ mean ± SD, n=10)^45,90^. Squash flowers are comparably large in size and volume to *Datura*, and provide habitats for commensal arthropods (Table S8)^91,92^. Squash flowers track ambient temperatures throughout the day, indicating they are passively heated by sunlight and are not endothermic (Fig. 1D, 1F). Despite this, they maintain consistently elevated relative humidity (ΔRH) during both open and closed phases, independent of ambient RH (Fig. 1E, 1G). This consistent elevation in ΔRH within squash flowers contrasts with the ephemeral humidity spike from nectar evaporation seen in the rapidly opening flowers of *Oenothera cespitosa* ^44^. The RH dynamics of *O. cespitosa* resemble the diel heat rhythms found in thermogenic flowers, fueled by the secretion of copious nectar rather than the combustion of starch reserves. In contrast, squash flowers exhibit a form of “hygroregulation”, akin to floral thermoregulation (persistent ΔT) documented for skunk cabbage (*Symplocarpus foetidus*), sacred lotus (*Nelumbo nucifera*), and various arum lilies ^85,86,93^.

Fine-scale measurements of T and RH gradients within squash flowers indicate that the nectary is often slightly warmer than ambient air during the flower’s open phase, coinciding with peak nectar secretion (Fig. 1h; 6:00 to 10:00; see Nepi 2001). This increase in floral T may result from the metabolic energy required for nectar secretion and/or the activity of microflora, such as yeast and bacteria, within the nectar ^94^. Zhang and Tang ^31^ show that floral temperature is generally higher in the center than at the periphery in several species of alpine plant communities suggesting that this phenomenon is likely more common. While corolla temperatures occasionally dip below ambient during the closed phase (Fig. 1I), ΔT fluctuations are minor compared to ΔRH. Humidity transects of *C. pepo* mirrored those of *Datura wrightii* ^45^, whereby humidity increases toward the nectary but dissipates at flower’s threshold (Fig. 2A, 2B). Flower manipulations revealed that petal transpiration is the major contributor to floral humidity, rather than nectar evaporation. The presence of stomata along the adaxial surface of squash corolla revealed patterns consistent with these humidity gradients (Fig. 2C).

While stomatal density per mm² on the adaxial surface is significantly lower in squash than in *D. wrightii* [see Fig. 1E in Dahake et al. 2022] ^45^, both species exhibit similar ΔRH levels. We hypothesize that the dense multicellular trichomes on squash petals likely enhance the boundary layer and facilitate permeability for water ^42,88^. Indeed, the water content of squash flowers is higher than that of *D. wrightii* (∼94% vs. ∼87% in *D. wrightii*). As with *D. wrightii*, squash flowers contain protoxylem that likely supports efficient water transport, maintaining turgidity in the floral hydrostatic skeleton (Fig. S1B) ^95^. Notably, the heightened humidity and distinct stomatal distribution patterns are also evident in the ancestral wild squash *Cucurbita foetidissima* in its native environment (Fig. 1A and Fig. 2D). This suggests that, while domestication has introduced substantial diversity in squash fruits, it has not fundamentally altered floral form and function. The similar microclimates between domesticated and wild squash flowers may thus support the co-expansion of squash and their specialist pollinators, *Xenoglossa pruinosa* ^47,48^.

Below we discuss the effects of squash floral microclimate on- (1) reproductive success, (2) behaviors of mutualists and antagonists, and (3) their functional roles as signals and cues for their visitors.

#### 1. Plant reproductive success

The unique microclimate within squash flowers has significant implications for pollen behavior, directly influencing male plant fitness. Prior studies have shown that humidity, temperature, and radiation (gamma and UV) can affect pollen germination rates in cucurbits and other flowering plants ^96–100^. Squash pollen is recalcitrant, meaning it is susceptible to desiccation damage ^101^, and our findings support this vulnerability.

Germination rates dropped precipitously within 3-4 hours post-anthesis, consistent with previous findings for cucurbit pollen ^102,103^. High levels of pollen rupture in humidity-reduced flowers highlight the critical role of humid floral microclimates in maintaining pollen viability. The persistence of high humidity even after flower senescence supports the hypothesis that flower closure or nyctinasty evolved to protect floral organs from environmental stress ^15,104^. The observation that male flowers have higher stomatal densities than female flowers suggest adaptive pressures to safeguard pollen integrity. Experiments on female flowers indicated that floral humidity manipulation also affected seed set and fruit quality in *Cucurbita pepo* (Fig. S4), though similar experiments are needed in wild squash, as domesticated varieties produce at least small fruits even without successful pollination. Beyond floral adaptations, squash pollen itself has evolved mechanisms to avoid desiccation and protect against predation. The oil coating on squash pollen may prevent dehydration and provide a nutrient-rich resource for developing *X. pruinosa* larvae. In contrast, the spiky exine of cucurbit pollen deters generalist *Bombus* bees, as it can damage adult hindguts and increase larval mortality^105^.

#### 2. Effects on behaviors of mutualists and antagonists

Squash flowers attract diverse insect visitors (Video S3), including thrips, *Drosophila* flies, *Melissodes* and *Lasioglossum* spp. solitary bees, the specialist pollinator *X. pruinosa*, generalist honeybees (*Apis mellifera*) and bumblebees (*Bombus impatiens*), and herbivores like striped (*Acalymma vittatum*) and spotted cucurbit beetles (*Diabrotica undecimpunctata*) ^65,78,106^. Manipulating floral humidity affected only the landing frequency of *X. pruinosa* bees (Fig. 5B), not generalist pollinators (Figs. 5D & 5F). While “chamber damage” and “limb damage” treatments caused similar mechanical disruptions, only chamber-damaged flowers reduced landing by *X. pruinosa*, indicating that bees responded to reduced RH rather than damage volatiles. In field observations, *X. pruinosa* often hovered closely but avoided landing on humidity-reduced flowers, suggesting that they may perceive humidity just before landing, akin to honeybees and bumblebees avoiding flowers marked by hydrocarbons ^107–109^, or altered electric fields^110^. In comparison, generalist pollinators like bumblebees do not show strong innate preferences for humid flowers, but they are able to quickly learn associations between rewards and humidity ^111,112^. Barman et al ^113^ show that an experimental drought stress affected all except scent traits in the Styrian oil pumpkin flowers (*Cucurbita pepo* L. subsp. *pepo* var. *styriaca* Greb.) which had a negative impact on bumblebee visitation frequency in a flight cage assay. Neither squash specialists nor generalists showed significant changes in foraging duration on manipulated flowers (Fig. 5C, 5E, 5G), likely because, despite manipulation, flower RH remained above ambient levels (Fig. 4). In a context of abundant floral resources and high nectar secretion rates ^114–116^, the suboptimal conditions in damaged flowers may have minimal deterrent effects on generalists.

#### 3. Functional roles as signals or cues

Our results showing a reduction in landing frequency of squash bees due to lower floral humidity reveal an indirect impact on plant fitness. It is reported that male squash bees are ten times more common on squash flowers than females^79^ and that seven consecutive visits by male squash bees optimized fruit set in *C. pepo,* indicating that repeat visits are important in this system^79^. Unlike honeybees and bumblebees that return to their nests after foraging, male squash bees are solitary wanderers, using squash flowers not only as food, but also as potential mating sites and shelters. Our finding that reduced floral RH negatively affects squash bee residence in flowers further highlights their strong mutualistic relationship and the critical role of the floral microclimate. Therefore, it can be argued that floral humidity functions as a signal encouraging squash bees to land and reside in the flowers, benefiting the plants as well as the bees. Compared with squash bees, the influence of squash floral humidity on their generalist pollinators is neutral (Fig. 5d-g), such that the provision of RH information neither benefits nor harms either party. In contrast, the interaction between squash plants and cucurbit beetles is antagonistic. Beetles gather in the flowers to mate, feed on nectar and floral tissue ^64^, defecate inside the flowers, and can transmit the pathogen *Erwinia tracheiphila* ^67^, which can lead to severe wilting within days. Beetle-induced perforations in the floral structure also alter the microclimate (Fig. 4). In our field experiments, manipulated flowers attracted fewer beetles, suggesting that the physical damage itself reduced their residence (Fig. 6). Prior studies indicate that herbivory can alter floral volatiles and trigger bitter cucurbitacin production, deterring additional beetle damage ^63,65^. However, humidity manipulations had a less significant effect on beetles than on bees, potentially because beetles’ smaller size keeps them within the corolla’s boundary layer, where their preferred microhabitat is less affected by our relatively small perforations in flowers. Consequently, in this context, floral humidity functions as a cue used by the beetles to exploit the floral microhabitat at the expense of the plant’s fitness.

### Flowers as Domiciles or Shelters

Another important interaction involving floral microclimates is the use of flowers as shelters ^23^. Large-volume flowers serve as critical microhabitats for insects, offering protection from predators and desiccation—essential for insects given their vulnerability to water loss ^117–119^. Notably, many giant flowers have evolved to attract small pollinators capable of dwelling within them for extended periods ^120,121^. Examples include beetles inhabiting flowers like Waterlilies ^33^, Arum lilies ^122^, and cycad cones ^123^, as well as flies residing within chamber flowers such as *Aristolochia* ^124–126^, *Rafflesia* ^127^, and *Ceropegia*^128^. Similarly, nearly every *C. foetidissima* flower we sampled in the Sonoran Desert was inhabited by *Drosophila pseudoobscura* males and females engaged in courtship, feeding, or aggression, all within the same flower (Fig. 1A, Video S1). While beetles and flies frequently use flowers as shelters, examples involving bees are less common but notable. Male *Melissodes* bees, for instance, have been observed resting in bowl-shaped flowers such as Globemallows and California poppies by nature photographers. The darkly pigmented flowers of *Iris atropurpurea* (Iridaceae) form chambers mimicking shelters, attracting male bees (e.g., *Synhalonia spectabilis*, *Megachile sicula*) that subsequently pollinate the flowers ^129^. These flowers provide protection from rain and a warm refuge, enabling earlier mate-seeking behavior ^130^. Similarly, male squash bees (*X. pruinosa*) also use *C. pepo* flowers as day and night shelters ^54–57^. In the drier ambient conditions of natural squash habitats, high floral humidity likely helps these bees mitigate water loss, a major stressor for desert bees ^131^. Supporting this, our data show a positive correlation between squash bee occupancy and ambient dryness (Fig. 6f). A similar trend was observed for beetles (Fig. 6g), indicating flowers serve as effective shelters for smaller insects. As climate change drives increased vapor pressure deficits ^132,133^, we predict a growing reliance on floral microcosms by insects to mitigate desiccation stress ^134^. This trend may be exacerbated by increasing plant-pollinator asynchrony ^106^ and reduced warming tolerance in many insect species ^135^. Future studies on traits impacting plant-pollinator behavior should consider floral humidity as an important factor when testing hypotheses.

## Acknowledgements

We thank Matthew Barret from CALS Entomology for giving us *C. pepo* saplings, and John Putnam for greenhouse care. We also thank Steven G. Persaud for helping A.D in the field, and Judy Swan for conducting a writing workshop attended by A.D. We thank Avehi Singh for feedback on the manuscript. We thank Strom’s farm and bakery (https://strom.ca/) for welcoming the University of Guelph team to collect data on squash flowers. Parts of this research were funded through Cornell Sigma Xi and Cornell Neurobiology and Behavior dept. grants awarded to A.D. P.N.S was supported by Conselho Nacional de Desenvolvimento Científico e Tecnológico (CNPq) [201568/2017-9]. D.O. was funded by a doctorate scholarship from Fulbright Colombia-MINCIENCIAS (Colombian Minister of Science, Technology, and Innovation) and the Federal Capacity Fund multistate project from the National Institute of Food and Agriculture (NIFA), United States, Department of Agriculture [#NYC-139848]. This project is part of the Accelerating Green Plant Innovation for Environmental and Economic Benefit Cluster and is funded by the Canadian Ornamental Horticulture Alliance (COHA-ACHO) and by the Government of Canada under the Canadian Agricultural Partnership’s AgriScience Program (Project # ASC-15). P.K also acknowledges support from the Natural Sciences and Engineering Research Council (NSERC Individual Discovery Grant RGPIN-2018-0482).

## Author contributions

A.D. and R.A.R. designed the research conducted by the Cornell University team, P.K. and C.C designed the research conducted by the University of Guelph team; A.D., C.C., D.O., J.C., and P.N.S. collected data. A.D performed all the data analysis, data curation, and interpretation with inputs from R.A.R; D.O. helped A.D. with statistical analysis; A.D. drafted the manuscript, and all authors reviewed it.

## Methods

### 1. Floral humidity and temperature measurements of wild squash flowers in Arizona, USA

We set out to measure baselines of floral temperature and humidity in a wild, large-flowered squash species *Cucurbita foetidissima* Kunth ^68^ growing in its natural habitat, a monsoon-seasonal grassland near the Santa Rita Mountains, Arizona, USA. Baseline data were collected from male and female flowers on July 29, 2021, on a roadside near Rosemont Ranch on Highway 62, Saguarita, Santa Cruz County, AZ, USA [GPS: 31°46’36.8“N 110°43’23.9”W].

We sampled floral data along roadside transects for two one-hour sampling periods, initiated at 07:20 and 09:00 hrs, respectively, as flowers opened near sunrise and began closing by 09:45 hrs. Temperature (T) and relative humidity (RH) data were collected using two hand-held Omega HH314A Humidity-Temperature Meters. The probes were inserted into the chamber-like corollas and held for 20-30 sec before noting the readings (to 0.1 decimal places). Most flowers were occupied by several insects, whose abundance and identity were noted. The RH gradient and thermal data were digitized and analyzed at the Florida Canyon field station (Univ. of Arizona), to which subsamples of male and female flowers were transported in Ziploc bags, on ice, for measurements of floral volume and stomatal density.

#### i. Floral volume measurements

Floral volumes were measured for 7 female and 6 male cut flowers of *C. foetidissima* by setting the flower upright within a large funnel for support, then filling the fused corolla to the brim with tap water and decanting into a 100 ml glass graduated cylinder marked with 1 ml gradations. Similarly, volumes of male and female *C. pepo, C. moschata*, and *C. maxima* flowers were measured from the blooming plants at squash farms near Cornell University, Ithaca, NY USA.

#### ii. Floral stomatal measurements

Floral stomata were counted by staining cuticular peels of the corolla surface for microscopy. Floral peels were taken at four locations from the adaxial surface of the corolla (see Fig. 2c inset). Location one was closest to the corolla base near the nectary. Subsequent peels were taken ∼1-2cm apart up to the apex of the corolla. Peels were stained with 50:50 Safranin: water solution for 15 seconds, subsequently rinsed with water, and mounted on microscope slides with coverslips. Peels were imaged at 10x under a Nikon Eclipse 80i microscope connected to a digital camera. All stomata within a 1mm^2^ area were counted using Image J.

### 2. Floral humidity and temperature measurements of cultivated squash flowers in Ontario, Canada

#### i. Micrometeorology of squash flowers across 24-hours

We performed baseline measurements of floral T and RH from commercially grown *Cucurbita pepo* in Ontario, Canada (Strom’s Farm https://strom.ca/, Guelph, ON; GPS: 43°29’52.4“N 80°17’32.3”W) in July 2018 and 2020. Flowers of *C. pepo* bloom from July to October at this site, with the floral pollination window spanning from mid-July to mid-August for fruits to be developed for commercial harvest in October. During this time, flowers generally open at sunrise and close at midday. To measure the RH and T of the floral microclimate throughout the transition to anthesis, nectar production and closing/wilting stages, four T and RH digital sensors (HygroVUE10, Campbell Scientific) were inserted into four male flowers. These sensors were connected to a CR1000X Campbell Scientific datalogger. The CR1000X has a standard operating range of - 40° to +70° and is powered by an external 12V battery. The sensors are required to be shielded from direct solar radiation for accurate data collection, so a shade net was set up above the canopy. The shading fabric was secured to the ground with wooden stakes at 1m height.

Floral buds were identified one day prior to anthesis and were located within a 2m radius of each other. The sensors were secured to metal plant stakes using plastic twist ties and inserted into the closed flower bud the night before floral opening. We left these sensors in place throughout floral opening and ensured that they did not contact the floral organs.

Ambient T and RH values were recorded by two HygroVUE10 sensors placed between focal flowers to evaluate whether metabolic heat production occurred during the enzymatic activity of nectar production. An InfraRed thermal camera (Teledyne FLIR) was used to take infrared images of both male and female *C. pepo* flowers at anthesis (06:00 hrs), noting the surface T of the floral organs during the period of floral opening and nectar production. Loggernet software was used to program and configure data collection.

#### ii. Focal comparisons of floral temperature dynamics with digital thermometers

Because many flowers are either thermogenic or reflect solar radiation (Van der Kooi et al. 2019), we explored whether the unusually large, chamber-like flowers of *Cucurbita pepo* might accumulate heat during anthesis. Accordingly, we performed targeted studies of floral temperature as flower buds of *C. pepo* matured, opened, and closed, across a range of flower sizes and under direct sunlit vs. shaded conditions. These measurements were taken at Strom’s Farm (field 1, as described above), where over 30 varieties of orange pumpkin are grown, and at a second site (field 2) in Peterborough, Ontario, Canada [GPS: 44°25’16.71“N, 78°20’46.57”W] in August 2018, at which we used flowers from the ‘Pik-a-pie’ (pie pumpkin) and ‘Spaghetti Squash’ (spaghetti squash) commercial varieties.

Ambient T (outside of the flower) and floral T (within the corolla tubes) were measured without touching the petals or sexual organs using copper-constantan thermocouples (type K, resolution 0.1C, accuracy 0.4% + 1C, Omega^TM^) and a 4-channel Data Logger Thermometer (RDXL 4SD, Omega^TM^). We noted whether all focal flowers were male (staminate) or female (pistillate) and blooming in the sun or shade during measurements. In addition to floral tube and ambient air T, we analyzed the differences between these measurements (hereafter, ΔT), which were positive when the floral T was warmer than ambient T and negative when the floral T was colder.

Given that ambient temperatures change during the morning of anthesis, we explored whether floral T changes with time of day. Specifically, we measured ΔT for whole flowers, corollas and nectary disks across 4-5 arbitrary time periods beginning at anthesis and ending with flower wilting, over 3 replicate days (July 27, 28, 30, 2020). We marked 6 male and 4 female flowers as they opened at dawn and performed repeated measures on them during all time periods until flowers wilted. T measurements for corolla and nectary disk were performed by contacting the surfaces of these floral tissues with the copper constantan thermocouple, whereas whole flower T was measured in the headspace within each flower, as described above.

#### iii. Continuous tracking of wilted flower chamber temperatures

Male squash bees are known to use the floral chambers of wilted, senescing squash flowers as domiciles when they are quiescent (see introduction). We sought to determine whether wilted flowers heat passively and might thereby provide warmth as well as shelter to male squash bees. Temperatures within and outside of aged, wilted flowers at field 2 of *C. pepo* were measured by copper-constantan thermocouples (type K, resolution 0.1C, accuracy 0.4% ± 1C, Omega^TM^) and a Multi-Channel Universal Input Touch Screen Data Logger (OM-DAQXL1, Omega^TM^), and were recorded at a rate of 12 samples/hour. In each trial we tracked seven wilted flowers, with a thermocouple inserted within each one. These flowers wilted two to three hours before the trial, and we avoided flowers that had wilted on the previous day. External to the focal wilted flowers, a thermocouple was secured in the shade at a similar height to the inside thermocouples to measure ambient air T. Trials were performed overnight in field 2 from 15:00 hrs to 05:00 hrs, three times for the Pik-a-Pie variety and two times for Spaghetti Squash (Data presented together). We calculated floral ΔT for each monitored hour by subtracting ambient air temperature from floral temperature.

### 3. Greenhouse cultivation of *C. pepo* for RH gradients and pollen viability experiments

*Cucurbita pepo* (Golden Zucchini) plants were grown from seeds (W. Atlee Burpee & Co and High Mowing Organic Seeds.) in summer of 2020 and 2021 in the rooftop greenhouses of Corson Mudd Hall, Cornell University, Ithaca NY, USA. The greenhouse was controlled from overheating with automatic exhaust fan and the artificial lighting period was set to 16:8 hours of light:dark cycle. Seeds were sown in a starter tray using 3:1 Lambert: perlite soil mixture (Lambert 111). Saplings were transplanted into half-gallon pots and watered daily with 21-5-20 fertilizer (nitrogen-phosphate-potash). Plants continued flowering for 4-6 weeks.

#### i. Vertical RH gradients of control and manipulated flowers

We used an Omega HH314A Humidity-Temperature Meter and a modified KD Scientific syringe pump to measure the baseline vertical RH gradients in the headspace of the newly opened male and female squash flowers ^also see 44,45^. A metal rod (0.5 cm diam) was screwed to the syringe pump at the end of which the humidity probe was attached. The hygrometer was inserted into the corolla tube making sure the anther and stigma filaments were intact. RH and T were recorded every second from the base of the corolla with the syringe pump moving vertically at 0.2 mm/sec ^see details in 45^ accounting for a total 14 cm vertical transect (Fig. 2a). A second humidity meter measured T and RH concurrently outside of the flower as ambient controls.

To determine the extent to which floral nectar or anatomical features (e.g. stomata or leaky cuticles in the corollas) are responsible for RH gradients in squash flowers, we conducted sequential experiments on the same set of flowers in the following way. In the first set, the vertical RH gradient was measured on the control (unmanipulated) flowers. In the second set, nectar was extracted with a 1ml syringe before taking RH measurements from the same flowers, testing the hypothesis that nectar evaporation contributes to high floral humidity during the morning hours after anthesis, as previously demonstrated for *Oenothera cespitosa* flowers ^44^. In the third set, a smear of petroleum jelly (Vaseline) was applied on the inner (adaxial) surface of the flower from the corolla base near the nectary to where the petals fuse (floral chamber), testing the alternative hypothesis that cuticular leakage and/or stomatal transpiration contribute to steep floral humidity gradients, as previously demonstrated for *Datura wrightii* flowers ^45^. For the Vaseline treatment, flowers were fanned to remove residual humid air before carrying out measurements. To calculate the difference in humidity (ΔRH), ambient RH values were subtracted from floral RH.

#### ii. Male fitness consequences of experimental manipulation of floral humidity

Given that male flowers were found to show a higher stomatal density than female flowers, we hypothesized that floral humidity is critical for pollen viability. Accordingly, we tested whether differences in floral RH impact the male fitness of squash flowers, using pollen rupture, germination, and ungerminated pollen as response variables. Pollen was collected from flowers exposed to three experimental treatments: control, Vaseline smeared, and excised corolla. In the control group, flowers were left untreated. We implemented the remaining two treatments to reduce floral RH by varying degrees. In the Vaseline-treated flowers, a cotton swab was used to cover the inner (adaxial) surface of the corolla with a thin layer of petroleum jelly. In the excised corolla group, the corolla was cut with scissors, exposing the stamens to ambient air without damaging them. The flowers were treated in the morning before the first round of pollen collection and left that way for the entire duration of the experiment. Three large Petri dishes labeled with treatment were prepared in the morning before pollen collection and floral manipulations. The bottom of each dish was lined with a moist Kimwipe to maintain a humid environment, then the dishes were carried to the greenhouse where the plants were located. Immediately before pollen collection, round microscope slide coverslips were placed within the Petri dishes (one coverslip per flower) and two drops of the germination solution were gently placed on each coverslip using a Pasteur pipette.

The pollen germination solution was prepared according to Brewbaker and Kwack ^69^. The recipe consisted of 10% sucrose, 100 ppm H_3_BO_3_, 300 ppm Ca (NO_3_)_2_·4H_2_O, 200 ppm MgSO_4_·7H_2_O, and 100 ppm KNO_3_ dissolved in distilled water. After pilot experiments, the concentration of Ca (NO_3_)_2_·4H_2_O was raised from 300 ppm to 800 ppm to improve the germination rates and reduce pollen tube entanglement.

Each treatment group replicate consisted of 3-5 plants, depending on how many *C. pepo* plants were blooming at a given time, in most cases using one flower per plant for pollen collection. Pollen collection was conducted using a pipette tip (20-200 μL) which was rubbed against a piece of polyester cloth to generate a static charge. The charged tip was then held immediately above the stamen of the flower to attract the pollen. This method reliably attracted 38.7±0.82 (mean±SEM, n= 378 collections) grains from each flower. The pipette tip was cleaned between each collection to remove pollen from other flowers. The collected pollen was then gently deposited on top of the coverslip containing the germination solution within its respective petri dish and a lid was placed on each petri dish.

The humidity of the flowers and surrounding environment for each treatment group was measured using an Omega HH314A Humidity Temperature Meter immediately after each pollen collection. The floral humidity was measured by holding the tip of the probe next to the stamen for 30 seconds and the surrounding humidity was measured by holding the probe 10-20 cm away from the treatment group of plants for 30 seconds. The collected pollen within the Petri dish was moved to the laboratory to germinate for approximately 45 minutes. The first pollen collection began at 07:30 hrs and the above procedures were repeated every hour until 12:30 hrs.

After the pollen was allowed to germinate for approximately 45 minutes after each collection, the Petri dishes with the germinated pollen on coverslips were then moved directly beneath a Nikon Eclipse 80i microscope. Each coverslip in the Petri dish was photographed with 10x magnification using the Amscope MU1000 microscope camera attached to the Nikon Eclipse 80i microscope. For each slide, the region with the highest pollen density was photographed (areas where the pollen clumped together were avoided since they obscured the view). Every pollen grain in the image except the ones touching the edge, or those that were completely out of focus, was counted for the total pollen count using the cell counter tool in Image J. For the viable pollen count, pollen was considered viable if the pollen germinated with pollen tube attached to the pollen grain, and the length of the pollen tube greater than the radius of the pollen grain. In the case that the pollen tube was detached from the pollen grain and floating debris was visible, such pollen was categorized as “germinated but ruptured”. In some images, none of the pollen tubes were visible but abundant debris around the pollen was evident. For such images, if the pollen appeared partially transparent, as an indication of the pollen material being expelled, it was considered “germinated but ruptured”. Pollen that looked completely opaque and intact with no sign of germination or pollen debris was considered “Ungerminable pollen” ^70^ (see images in Fig. 3).

### 4. Field observations of insect visits, foraging time, and residence in experimentally manipulated squash flowers

We explored the potential impact of floral micrometeorology on insect visitors and residents of squash flowers. Field observations of pollinator visits to squash flowers were conducted for two consecutive years at Cornell University’s Agricultural Experimental Station ^71^ managed by the College of Agricultural and Life Sciences (CALS). Field observations lasted from July 27-Aug 16, 2021 (9 days) at the Maple Avenue [GPS coordinates: 42.441564, -76.472407] farm plot, and from July 15-Aug 20, 2022 (10 days), split between Maple Avenue [GPS: 42.441079, -76.471109] and Freeville farm [GPS: 42.521037, -76.328198] plots. The farm plots were designed to grow *C. pepo* (Golden zucchini), *C. maxima* (103 & 105 NOVIC), and *C. moschata* (Bugle x Dickinson) varieties of squash. Seeds were sown during the third week of May in starter trays with Cornell Soil-less potting mix. After two weeks, samplings were transplanted 2ft apart in the farm plots using a water wheel transplanter. The plots had Sandy Loam or Gravely Loam soil type which was amended with 13-13-13 NPK and plastic mulch was laid on the raised bed. At least 1.5 m distance was maintained between two raised beds. All plots used organic soil and were free of pesticides. Plots were irrigated weekly using water from nearby creeks.

To manipulate floral humidity in the field, we introduced perforations either in the flower chamber to allow humidity to escape or in the flower limb as a sham control for the mechanical damage to the corolla, measuring floral T and RH gradients as described above. This experimental setup was inspired by observations of widespread damage caused by cucurbit beetles attacking flowers on squash farms. While beetle herbivory predominantly occurred on the flower limb, instances of damage within the floral chamber were also noted (Fig. 4b). Given the dynamics of the vertical gradient of humidity, we predicted that these two modes of damage would have distinct effects on floral micrometeorology. Limb damage was expected to have minimal impact on the microenvironment within the floral chamber, whereas chamber damage could disrupt humidity and temperature levels.

For the purposes of suitable replication, we limited observations to male flowers, due to their greater abundance (10:1 male: female flower ratio) ^72^. In each triad of flowers, two were experimentally manipulated and the third was left as an undamaged control flower. In the “chamber-damage” treatment, a 2.5 cm diameter hole was punched at the bottom half of the flower chamber to allow floral humidity to leak out (see Fig. 4b for images). For the “limb-damage” treatment, a hole was punched on the top half of the flower on the corolla limb as a sham control for the damage caused to the flowers in the “chamber-damage” treatment. The “limb damage” hole does not leak humidity from the floral chamber but accounts for wound volatiles and the physiological changes associated with damage to the flowers. The third flower was left undamaged as true control but was handled in the same way as the other two flowers. Immediately after the holes were punched (using a steel leather hole puncher), the edges of the holes were sealed with petroleum jelly to prevent water loss.

In the first experiment, we focused on evaluating the impact on bees (both male and female) as floral visitors. Given that squash flowers attract both generalist and specialist bees as pollinators, we anticipated that the altered floral microenvironment would influence their foraging behavior in distinct ways, such as visitation frequency and foraging duration.

We monitored the visitation frequency of bee visitors to squash flowers with and without perforations over three periods in the morning hours when the flowers were typically open. Initially, we examined whether there were disparities in bee visitation rates across the various flower treatments, encompassing all observed bee species.

For the pollinator visitation experiments in 2021, observations were conducted by one or two people on 3-5 plants per day for three consecutive observation periods during morning hours. Period 1 was conducted from 07:00–08:30 hrs, period 2 from 08:30-09:30 hrs, and period 3 from 10:00-11:00 hrs. Plants were selected and tagged before the first observation period. Only those plants that had three blooming flowers within ∼100 cm distance of each other were selected, because the observer needed to score insect visits to all flowers simultaneously.

For each observation, the observer typically stood 1 m away from the flowers with a pen and a datasheet, set a 10-minute stopwatch on their phone, and scored the entry and exit times of all insects visiting any of the three flowers. We considered a “visit” only if the insect visitor entered the floral chamber or landed on the corolla. Close approaches or hovers near the floral surface were not considered visits and therefore were not scored. After 10 min, each observer recorded the ambient and floral T and RH before moving on to the next tagged plant patch. For the second and third periods, the observer returned to the same plants tagged in period 1 and repeated the process.

The data were analyzed for two response variables: visitation frequency and the duration of each visit (foraging duration). The visitation frequency is defined as the number of independent landings noted for each visitor type, such as bumblebees, squash bees, or honeybees. Higher visitation frequencies are often attributed to greater outcrossing benefits to the plants ^73–77^, increasing fruit yield and seed set ^78,79^. The foraging duration was calculated as the difference in time between landing and exiting the flower by each visitor. The foraging duration refers to the time spent in the flower seeking nectar and pollen and often attributed to greater pollen loading on the bodies of insects or greater deposition on the stigma due to moving around in the floral chamber of female flowers ^78,80^. Pollen loading by female *X. pruinosa* bees can also influence the reproductive success of the bees since they provision larvae with squash pollen for nutrition ^80^.

In the second experiment, we tested whether floral manipulations and their impacts on RH affect specialized insects that use faded, closed flowers as shelter. These floral residents include the specialist cucurbit beetle herbivores and male squash bees as specialist pollinators. Field scoring of floral residents was conducted July-Aug 2022 on two separate farm plots (Maple Avenue and Freeville) in Tompkins Co., NY, USA. We made 10 total visits and pooled data from both farms. Floral manipulations were performed between 09:00-10:00 hrs as described above for 10-19 plants with three flowers each and were tagged with florescent ribbons to keep track of the treatments. The closed and wilted (male) squash flowers were revisited the same day between 14:00-15:00 hrs and were gently opened by the observer to count the number of insects residing within the flower chambers. The squash bees were easily disturbed and quickly flew out of the flower chamber. Most flowers contained 1-2 bees, however some flowers housed up to 4 bees. The beetles were counted by gently exhaling on them and narrowing the flower opening such that they could only exit one at a time. This method was useful to avoid miscounting, especially since some flowers were occupied by up to 60 beetles (Video S2). The count data were immediately scored on a pad and digitized in the lab, which was followed by measurements of floral RH and temperature for flowers of all treatments. Due to hygrosensor malfunction and a couple of rainy days, we were able to note floral RH and T data for only 4 out of the 10 experimental visits to the farms in 2022. Ambient T and RH values were recorded on most visits or obtained from local weather stations. For vapor pressure deficits (VPD), we first calculated saturation vapor pressure (SVP) with the formula: 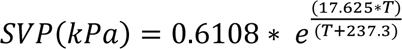 where *T* is the temperature in degrees Celsius, and e is the base of the natural logarithm. Absolute vapor pressure (AVP) was calculated with the formula: 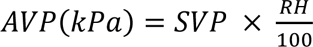 and vapor pressure deficit was calculated with the formula: *VPD*(*kPa*) = *SVP* − *AVP*

### 5. Statistics

All statistics and model fits were performed in R v4.3.2 ^81^. To evaluate whether *C. foetidissima* floral ΔT and ΔRH are above ambient levels (Fig. 1b and c), we performed a One-sample t-test against zero. The temperature of different floral parts (Fig. 1h) was analyzed using a linear mixed-effects model with temperature values log transformed to approach normal distribution for the residuals. In the model, temperature served as the response variable and floral parts interaction with sex as the fixed effect independent variables. We initially added random effects of flower ID and day to the model, but flower ID contributed zero variance and hence was dropped from our final model.

We fitted the vertical gradients of the floral humidity shown in Fig. 2a and 2b with a nonlinear exponential decay model of form 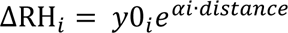 where *y0* is the intercept, *α* is the decay rate, and *i* is the treatment, using the nlme package ^82^. First, we fitted a single curve to the overall data to obtain starting values for y0 (39 for male and 37 for female flowers) and used 1 for decay rate. In subsequent models we allowed α and y0 to vary by treatment i and in the most complicated model we also included random effects of individual flowers on α and y0. In all cases, the most complicated model was the best fit and lowest in AIC. We extracted model estimates and confidence intervals for pairwise comparisons between treatments using the emmeans package ^83^ to determine if the floral treatments had a significant effect on the slope or intercept of the vertical humidity gradients. P-values were adjusted for multiple comparisons.

Pollen viability data were square root transformed before performing a two-way ANOVA with time + treatment as the explanatory variable and pollen behavior as the response variable. For pairwise comparisons between treatments, we performed a post hoc Tukey’s test for subsets of the data for each timepoint and adjusted the p-values for multiple comparisons.

We used a generalized linear mixed effect model (GLMM) with Poisson distribution to analyze bee visitation frequency. Counts of individual bees served as the response variable, treatment * period was the fixed effect and plant variety, day, and site were included as random effects in the initial model and dropped in subsequent models if they did not contribute to the variance of the model. For honeybee visitation frequency we used a zero-inflated GLM with Poisson log link due to the fewer counts of honeybees. For those GLMs where the treatment had a significant impact on visitation rates, we conducted pairwise comparisons within a period using “emmeans” package and adjusted p-values with Tukey-Kramer method.

We used a linear mixed-effects model (LMM) to analyze the foraging duration of each visitor. In the model, foraging time of individual bees was the response variable, treatment* period was the fixed effect explanatory variable, and plant variety, day, and site were included as random effects in the initial model but dropped in subsequent models if they did not contribute to the variance of the model.

We analyzed beetle residence using a generalized linear model (GLM) with Poisson distribution. Beetle count was the response variable and treatment, plant variety and site were the explanatory variables. The squash bee residence was analyzed using a zero-inflated GLM with Poisson distribution since only ∼11% of the flowers were occupied by the bees.

Correlations between vapor pressure deficit (VPD) and beetle counts in flowers were analyzed with a Pearson’s product-moment correlation test and plotted with the ggscatter function in R. The squash bees were analyzed similarly, but because bees were not as abundant as beetles in flowers, we performed a correlation between VPD and the proportion of flowers that were occupied by bees, instead of the absolute number of bees, for each of the 10 observation days.

## Supplemental information

Document S1. Figures S1–S4 and Tables S1 and S8

Video S1. *Cucurbita foetidissima* floral chamber occupied by mating and courting *Drosophila pseudoobscura*, related to Figure 1

Video S1 *Cucurbita pepo* flowers inhabited by tens of cucurbit beetles and other insects, related to figure 6

## References

1. Harder, L.D., and Johnson, S.D. (2009). Darwin’s beautiful contrivances: evolutionary and functional evidence for floral adaptation. New Phytol. 183, 530–545. 10.1111/j.1469-8137.2009.02914.x.

2. Schiestl, F.P., and Johnson, S.D. (2013). Pollinator-mediated evolution of floral signals. Trends Ecol. Evol. 28, 307–315. 10.1016/j.tree.2013.01.019.

3. Trunschke, J., Sletvold, N., and Ågren, J. (2017). Interaction intensity and pollinator-mediated selection. New Phytol. 214, 1381–1389. 10.1111/nph.14479.

4. Wright, G.A., and Schiestl, F.P. (2009). The evolution of floral scent: the influence of olfactory learning by insect pollinators on the honest signalling of floral rewards. Funct. Ecol. 23, 841–851. 10.1111/j.1365-2435.2009.01627.x.

5. Koski, M.H. (2020). The role of sensory drive in floral evolution. New Phytol. 227, 1012–1024. 10.1111/nph.16510.

6. Raguso, R.A. (2004). Flowers as sensory billboards: progress towards an integrated understanding of floral advertisement. Curr. Opin. Plant Biol. 7, 434–440. 10.1016/j.pbi.2004.05.010.

7. Strauss, S.Y., and Whittall, J.B. (2006). Non-pollinator agents of selection on floral traits. In Ecology and Evolution of Flowers (Oxford University PressOxford), pp. 120–138. 10.1093/oso/9780198570851.003.0007.

8. Caruso, C.M., Eisen, K.E., Martin, R.A., and Sletvold, N. (2019). A meta-analysis of the agents of selection on floral traits. Evolution 73, 4–14. 10.1111/evo.13639.

9. Knauer, A.C., Kokko, H., and Schiestl, F.P. (2021). Pollinator behaviour and resource limitation maintain honest floral signalling. Funct. Ecol. 35, 2536–2549. 10.1111/1365-2435.13905.

10. Armbruster, W.S., Lee, J., and Baldwin, B.G. (2009). Macroevolutionary patterns of defense and pollination in *Dalechampia* vines: adaptation, exaptation, and evolutionary novelty. Proc. Natl. Acad. Sci. U. S. A. 106, 18085–18090. 10.1073/pnas.0907051106.

11. Chautá, A., Kumar, A., Mejia, J., Stashenko, E.E., and Kessler, A. (2022). Defensive functions and potential ecological conflicts of floral stickiness. Sci. Rep. 12, 19848. 10.1038/s41598-022-23261-2.

12. Frey, F.M. (2004). Opposing natural selection from herbivores and pathogens may maintain floral-color variation in *Claytonia virginica* (Portulacaceae). Evolution 58, 2426–2437. 10.1111/j.0014-3820.2004.tb00872.x.

13. Kessler, D., Gase, K., and Baldwin, I.T. (2008). Field experiments with transformed plants reveal the sense of floral scents. Science 321, 1200–1202. 10.1126/science.1160072.

14. Huang, M., Sanchez-Moreiras, A.M., Abel, C., Sohrabi, R., Lee, S., Gershenzon, J., and Tholl, D. (2012). The major volatile organic compound emitted from *Arabidopsis thaliana* flowers, the sesquiterpene (E)-β-caryophyllene, is a defense against a bacterial pathogen. New Phytol. 193, 997–1008. 10.1111/j.1469-8137.2011.04001.x.

15. Sun, J.-F., Gong, Y.-B., Renner, S.S., and Huang, S.-Q. (2008). Multifunctional bracts in the dove tree *Davidia involucrata* (Nyssaceae: Cornales): rain protection and pollinator attraction. Am. Nat. 171, 119–124. 10.1086/523953.

16. Koski, M.H., and Ashman, T.-L. (2015). Floral pigmentation patterns provide an example of Gloger’s rule in plants. Nat. Plants 1, 14007. 10.1038/nplants.2014.7.

17. Kevan, P.G. (1975). Sun-tracking solar furnaces in high arctic flowers: significance for pollination and insects. Science 189, 723–726. 10.1126/science.189.4204.723.

18. Galen, C., and Stanton, M.L. (2003). Sunny-side up: flower heliotropism as a source of parental environmental effects on pollen quality and performance in the snow buttercup, *Ranunculus adoneus* (Ranunculaceae). Am. J. Bot. 90, 724–729. 10.3732/ajb.90.5.724.

19. Li, J.-K., and Huang, S.-Q. (2009). Flower thermoregulation facilitates fertilization in Asian sacred lotus. Ann. Bot. 103, 1159–1163. 10.1093/aob/mcp051.

20. Seymour, R.S., Ito, Y., Onda, Y., and Ito, K. (2009). Effects of floral thermogenesis on pollen function in Asian skunk cabbage *Symplocarpus renifolius*. Biol. Lett. 5, 568–570. 10.1098/rsbl.2009.0064.

21. Vannette, R.L. (2020). The floral microbiome: Plant, pollinator, and microbial perspectives. Annu. Rev. Ecol. Evol. Syst. 51, 363–386. 10.1146/annurev-ecolsys-011720-013401.

22. Fetters, A.M., and Ashman, T.-L. (2023). The pollen virome: A review of pollen-associated viruses and consequences for plants and their interactions with pollinators. Am. J. Bot. 110, e16144. 10.1002/ajb2.16144.

23. Raguso, R.A. (2023). Hidden worlds within flowers. Curr. Biol. 33, R506–R512. 10.1016/j.cub.2023.04.054.

24. Kevan, P.G., Tikhmenev, E.A., and Nunes-Silva, P. (2019). Temperatures within flowers and stems: Possible roles in plant reproduction in the north. Vestn. Severo. Vost. Nauchnogo Tsentra Dalnevost. Otd. RAN, 38–47. 10.34078/1814-0998-2019-1-38-47.

25. Coates, C., and Kevan, P. (2021). Exploring Micrometeorology in Plant Stems and Flowers. Scientia 10.33548/SCIENTIA666. https://doi.org/10.33548/SCIENTIA666.

26. Kevan, P.G. (1970). High arctic insect-flower relations : the inter-relationships of arthropods and flowers at Lake Hazen, Ellesmere Island, N.W.T., Canada.

27. Kevan, P.G. (2020). Heat accumulation in hollow Arctic flowers: possible microgreenhouse effects in syncalyces of campions (*Silene* spp. (Caryophyllaceae)) and zygomorphic sympetalous corollas of louseworts (*Pedicularis* spp. (Orobanchaceae)). Polar Biology 43, 2101–2109. 10.1007/s00300-020-02772-6.

28. van der Kooi, C.J., Kevan, P.G., and Koski, M.H. (2019). The thermal ecology of flowers. Ann. Bot. 124, 343–353. 10.1093/aob/mcz073.

29. Kevan, P.G., and Coates, C. (2024). Heliocaminiform structures: plant organs that function as microgreenhouses. FACETS 9, 1–20. 10.1139/facets-2023-0099.

30. Harrap, M.J., Rands, S.A., Hempel de Ibarra, N., and Whitney, H.M. (2017). The diversity of floral temperature patterns, and their use by pollinators. Elife 6, e31262. 10.7554/eLife.31262.

31. Zhang, Y., and Tang, Y. (2023). Flower surface is warmer in center than at edges in alpine plants: evidence from Qinghai-Tibetan Plateau. J. Plant Ecol. 16. 10.1093/jpe/rtad023.

32. Raskin, I., Turner, I.M., and Melander, W.R. (1989). Regulation of heat production in the inflorescences of an *Arum* lily by endogenous salicylic acid. Proc. Natl. Acad. Sci. U. S. A. 86, 2214–2218. 10.1073/pnas.86.7.2214.

33. Seymour, R.S., and Matthews, P.G.D. (2006). The role of thermogenesis in the pollination biology of the Amazon waterlily *Victoria amazonica*. Ann. Bot. 98, 1129– 1135. 10.1093/aob/mcl201.

34. Körner, C., and Hiltbrunner, E. (2018). The 90 ways to describe plant temperature. Perspect. Plant Ecol. Evol. Syst. 30, 16–21. 10.1016/j.ppees.2017.04.004.

35. Peris, D., Postigo-Mijarra, J.M., Peñalver, E., Pellicer, J., Labandeira, C.C., Peña-Kairath, C., Pérez-Lorenzo, I., Sauquet, H., Delclòs, X., and Barrón, E. (2024). The impact of thermogenesis on the origin of insect pollination. Nat. Plants 10, 1297– 1303. 10.1038/s41477-024-01775-z.

36. Thien, L.B., Azuma, H., and Kawano, S. (2000). New perspectives on the pollination biology of basal angiosperms. Int. J. Plant Sci. 161, S225–S235. 10.1086/317575.

37. Seymour, R.S., White, C.R., and Gibernau, M. (2003). Heat reward for insect pollinators. Nature 426, 243–244. 10.1038/426243a.

38. Terry, L.I., Roemer, R.B., Booth, D.T., Moore, C.J., and Walter, G.H. (2016). Thermogenic respiratory processes drive the exponential increase of volatile organic compound emissions in *Macrozamia* cycad cones. Plant Cell Environ. 39, 1588– 1600. 10.1111/pce.12730.

39. Barthlott, W., Szarzynski, J., Vlek, P., Lobin, W., and Korotkova, N. (2009). A torch in the rain forest: thermogenesis of the Titan arum (*Amorphophallus titanum*). Plant Biol. (Stuttg.) 11, 499–505. 10.1111/j.1438-8677.2008.00147.x.

40. Angioy, A.M., Stensmyr, M.C., Urru, I., Puliafito, M., Collu, I., and Hansson, B.S. (2004). Function of the heater: the dead horse arum revisited. Proc. R. Soc. Edinb. Biol. 271 *Suppl 3*, S13–5. 10.1098/rsbl.2003.0111.

41. Galen, C. (2006). Solar furnaces or swamp coolers: Costs and benefits of water use by solar-tracking flowers of the alpine snow buttercup, *Ranunculus adoneus*. Oecologia 148, 195–201.

42. Roddy, A.B., Guilliams, C.M., Fine, P.V.A., Mambelli, S., Dawson, T.E., and Simonin, K.A. (2023). Flowers are leakier than leaves but cheaper to build. bioRxiv, 2076–2082. 10.1101/2023.04.11.536372.

43. Harrap, M.J.M., Hempel de Ibarra, N., Knowles, H.D., Whitney, H.M., and Rands, S.A. (2020). Floral humidity in flowering plants: A preliminary survey. Front. Plant Sci. 11, 249. 10.3389/fpls.2020.00249.

44. von Arx, M., Goyret, J., Davidowitz, G., and Raguso, R.A. (2012). Floral humidity as a reliable sensory cue for profitability assessment by nectar-foraging hawkmoths. Proc. Natl. Acad. Sci. U. S. A. 109, 9471–9476. 10.1073/pnas.1121624109.

45. Dahake, A., Jain, P., Vogt, C.C., Kandalaft, W., Stroock, A.D., and Raguso, R.A. (2022). A signal-like role for floral humidity in a nocturnal pollination system. Nat. Commun. 13, 7773. 10.1038/s41467-022-35353-8.

46. Corbet, S.A., Unwin, D.M., and Prŷs-Jones, O.E. (1979). Humidity, nectar and insect visits to flowers, with special reference to *Crataegus, Tilia* and *Echium*. Ecol. Entomol. 4, 9–22. 10.1111/j.1365-2311.1979.tb00557.x.

47. López-Uribe, M.M., Cane, J.H., Minckley, R.L., and Danforth, B.N. (2016). Crop domestication facilitated rapid geographical expansion of a specialist pollinator, the squash bee *Peponapis pruinosa*. Proc. Biol. Sci. 283. 10.1098/rspb.2016.0443.

48. Pope, N.S., Singh, A., Childers, A.K., Kapheim, K.M., Evans, J.D., and López-Uribe, M.M. (2023). The expansion of agriculture has shaped the recent evolutionary history of a specialized squash pollinator. Proc. Natl. Acad. Sci. U. S. A. 120, e2208116120. 10.1073/pnas.2208116120.

49. Decker, D.S. (1988). Origin(s), evolution, and systematics of *Cucurbita pepo* (Cucurbitaceae). Econ. Bot. 42, 4–15. 10.1007/bf02859022.

50. Nass, U. (2024). USDA Economics, Statistics and Market Information. https://usda.library.cornell.edu/.

51. Artz, D.R., Hsu, C.L., and Nault, B.A. (2011). Influence of honey bee, *Apis mellifera*, hives and field size on foraging activity of native bee species in pumpkin fields. Environ. Entomol. 40, 1144–1158. 10.1603/EN10218.

52. Freitas, F.V., Branstetter, M.G., Franceschini-Santos, V.H., Dorchin, A., Wright, K.W., López-Uribe, M.M., Griswold, T., Silveira, F.A., and Almeida, E.A.B. (2023). UCE phylogenomics, biogeography, and classification of long-horned bees (Hymenoptera: Apidae: Eucerini), with insights on using specimens with extremely degraded DNA. Insect Syst. Divers. 7. 10.1093/isd/ixad012.

53. Hurd, P.D., Jr, and Gorton Linsley, E. (1964). The Squash and Gourd Bees— Genera Peponapis Robertson and Xenoglossa Smith— Inhabiting America North of Mexico (Hymenoptera: Apoidea). Hilgardia 35, 375.

54. Michener, C.D., and Lange, R.B. (1958). Observations on the ethology of neotropical Anthophorine bees (Hymenoptera: Apoidea). University of Kansas Science Bulletin 39, 69–96.

55. Hurd, P.D., Linsley, E.G., and Michelbacher, a. D. (1974). Ecology of the squash and gourd bee, Peponapis pruinosa, on cultivated cucurbits in California (Hymenoptera: Apoidea). Smithsonian Contributions to Zoology 168, 1–17. 10.5479/si.00810282.168.

56. Hurd, P.D., Linsley, E.G., and Thomas, W.W. (1971). Squash and gourd Bees (Peponapis, Xenoglossa) and the origin of the cultivated Cucurbita. Evolution 25, 218–234.

57. Willis, D.S., and Kevan, P.G. (1995). Foraging dynamics of *Peponapis pruinosa* (Hymenoptera: Anthophoridae) on pumpkin (*Cucurbita pepo*) in southern Ontario. Can. Entomol. 127, 167–175. 10.4039/ent127167-2.

58. Mathewson, J.A. (1968). Nest Construction and Life History of the Eastern Cucurbit Bee, *Peponapis pruinosa* (Hymenoptera: Apoidea). J. Kans. Entomol. Soc. 41, 255– 261.

59. Rozen, J.G., and Ayala, R. (1987). Nesting Biology of the Squash Bee *Peponapis utahensis* (Hymenoptera; Anthophoridae; Eucerini). J. N. Y. Entomol. Soc. 95, 28– 33.

60. Tepedino, V.J. (1981). The pollination efficiency of the squash bee (*Peponapis pruinosa*) and the honey bee (*Apis mellifera*) on summer squash (*Cucurbita pepo*). J. Kans. Entomol. Soc.

61. Vidal, M. das G., Jong, D. de, Wien, H.C., and Morse, R.A. (2010). Pollination and fruit set in pumpkin (*Cucurbita pepo*) by honey bees. Rev. Bras. Bot. 33, 106–113. 10.1590/s0100-84042010000100010.

62. Artz, D.R., and Nault, B.A. (2011). Performance of *Apis mellifera*, *Bombus impatiens*, and *Peponapis pruinosa* (Hymenoptera: Apidae) as pollinators of pumpkin. J. Econ. Entomol. 104, 1153–1161. 10.1603/ec10431.

63. Theis, N., Kesler, K., and Adler, L.S. (2009). Leaf herbivory increases floral fragrance in male but not female *Cucurbita pepo* subsp. *texana* (Cucurbitaceae) flowers. Am. J. Bot. 96, 897–903. 10.3732/ajb.0800300.

64. Theis, N., and Adler, L.S. (2012). Advertising to the enemy: enhanced floral fragrance increases beetle attraction and reduces plant reproduction. Ecology 93, 430–435. 10.1890/11-0825.1.

65. Theis, N., Barber, N.A., Gillespie, S.D., Hazzard, R.V., and Adler, L.S. (2014). Attracting mutualists and antagonists: plant trait variation explains the distribution of specialist floral herbivores and pollinators on crops and wild gourds. Am. J. Bot. 101, 1314–1322. 10.3732/ajb.1400171.

66. Andersen, J.F., and Metcalf, R.L. (1987). Factors influencing distribution of *Diabrotica* spp. in blossoms of cultivated *Cucurbita* spp. J. Chem. Ecol. 13, 681–699. 10.1007/BF01020152.

67. Sasu, M.A., Seidl-Adams, I., Wall, K., Winsor, J.A., and Stephenson, A.G. (2010). Floral transmission of *Erwinia tracheiphila* by cucumber beetles in a wild *Cucurbita pepo*. Environ. Entomol. 39, 140–148. 10.1603/EN09190.

68. Bemis, W.P., L. D. Curtis, C. W. Weber, and Berry, J. (1978). The Feral Buffalo Gourd, *Cucurbita foetidissima*. Econ. Bot. 32, 87–95.

69. Brewbaker, J.L., and Kwack, B.H. (1963). The essential role of calcium ion in pollen germination and pollen tube growth. Am. J. Bot. 50, 859. 10.2307/2439772.

70. Christensen, S.M., Munkres, I., and Vannette, R.L. (2021). Nectar bacteria stimulate pollen germination and bursting to enhance microbial fitness. Curr. Biol. 31, 4373–4380.e6. 10.1016/j.cub.2021.07.016.

71. CornellAES Cornell University Agricultural Experimental Station. https://cals.cornell.edu/agricultural-experiment-station/research-farms/campus-area-farms. https://cals.cornell.edu/agricultural-experiment-station/research-farms/campus-area-farms.

72. Whitaker, T.W. (1931). Sex ratio and sex expression in the cultivated cucurbits. Am. J. Bot. 18, 359. 10.2307/2435848.

73. Kalisz, S., Vogler, D.W., and Hanley, K.M. (2004). Context-dependent autonomous self-fertilization yields reproductive assurance and mixed mating. Nature 430, 884– 887. 10.1038/nature02776.

74. Brunet, J., and Sweet, H.R. (2006). Impact of insect pollinator group and floral display size on outcrossing rate. Evolution 60, 234–246. 10.1111/j.0014-3820.2006.tb01102.x.

75. Koski, M.H., Galloway, L.F., and Busch, J.W. (2019). Pollen limitation and autonomous selfing ability interact to shape variation in outcrossing rate across a species range. Am. J. Bot. 106, 1240–1247. 10.1002/ajb2.1342.

76. Duffy, K.J., Patrick, K.L., and Johnson, S.D. (2020). Outcrossing rates in a rare “ornithophilous” aloe are correlated with bee visitation. Plant Syst Evol 306. 10.1007/s00606-020-01656-w.

77. Christopher, D.A., Karron, J.D., Semski, W.R., Smallwood, P.A., Trapnell, D.W., and Mitchell, R.J. (2021). Selfing rates vary with floral display, pollinator visitation and plant density in natural populations of *Mimulus ringens*. J. Evol. Biol. 34, 803–815. 10.1111/jeb.13781.

78. McGrady, C.M., Troyer, R., and Fleischer, S.J. (2020). Wild bee visitation rates exceed pollination thresholds in commercial Cucurbita agroecosystems. J. Econ. Entomol. 113, 562–574. 10.1093/jee/toz295.

79. Cane, J.H., Sampson, B.J., and Miller, S.A. (2011). Pollination value of male bees: the specialist bee *Peponapis pruinosa* (Apidae) at cultivated summer squash (*Cucurbita pepo*). Environmental Entomology 40, 614.

80. Chan, S., and Raine, N.E. (2023). Sharing the wealth: pollen partitioning in a Cucurbita crop pollination system with reference to the wild hoary squash bee. J. Pollinat. Ecol. 33, 228–238. 10.26786/1920-7603(2023)751.

81. R Core Team (2023). R: A Language and Environment for Statistical Computing.

82. José C. Pinheiro and Douglas M. Bates (2024). nlme: Linear and Nonlinear Mixed Effects Models.

83. Lenth, R.V. (2024). emmeans: Estimated Marginal Means, aka Least-Squares Means.

84. Seymour, R.S., and Schultze-Motel, P. (1997). Heat-producing flowers. Endeavour 21, 125–129. 10.1016/s0160-9327(97)80222-0.

85. Seymour, R.S., and Schultze-Motel, P. (1998). Physiological temperature regulation by flowers of the sacred lotus. Philos. Trans. R. Soc. Lond. B Biol. Sci. 353, 935– 943. 10.1098/rstb.1998.0258.

86. Knutson, R.M. (1974). Heat production and temperature regulation in eastern skunk cabbage. Science 186, 746–747. 10.1126/science.186.4165.746.

87. Seymour, R.S. (2010). Scaling of heat production by thermogenic flowers: limits to floral size and maximum rate of respiration: Scaling of thermogenic flowers. Plant Cell Environ. 33, 1474–1485. 10.1111/j.1365-3040.2010.02190.x.

88. Roddy, A.B., Schreel, J.D.M., Paiva, D.C., Qin, N., Jiang, G.-F., Brodersen, C.R., and Simonin, K.A. (2024). Transpirational water loss from flowers is low but unregulated. bioRxiv, 2024.06. 28.600864. 10.1101/2024.06.28.600864.

89. Patiño, S., Grace, J., and Bänziger, H. (2000). Endothermy by flowers of *Rhizanthes lowii* (Rafflesiaceae). Oecologia 124, 149–155. 10.1007/s004420050001.

90. Dahake, A.S. (2023). The impact of floral and ambient humidity on the behavior and physiology of a nocturnal pollinator.

91. Raguso, R.A., Henzel, C., Buchmann, S.L., and Nabhan, G.P. (2003). Trumpet flowers of the sonoran desert: Floral biology of *Peniocereus* Cacti and Sacred *Datura*. Int. J. Plant Sci. 164, 877–892. 10.1086/378539.

92. Liu, C.-Q., Gao, Y.-D., Niu, Y., Xiong, Y.-Z., and Sun, H. (2019). Floral adaptations of two lilies: implications for the evolution and pollination ecology of huge trumpet-shaped flowers. Am. J. Bot. 106, 622–632. 10.1002/ajb2.1275.

93. Wagner, A.M., Krab, K., Wagner, M.J., and Moore, A.L. (2008). Regulation of thermogenesis in flowering Araceae: the role of the alternative oxidase. Biochim. Biophys. Acta 1777, 993–1000. 10.1016/j.bbabio.2008.04.001.

94. Herrera, C.M., and Pozo, M.I. (2010). Nectar yeasts warm the flowers of a winter-blooming plant. Proc. Biol. Sci. 277, 1827–1834. 10.1098/rspb.2009.2252.

95. Roddy, A.B., Simonin, K.A., McCulloh, K.A., Brodersen, C.R., and Dawson, T.E. (2018). Water relations of *Calycanthus* flowers: Hydraulic conductance, capacitance, and embolism resistance. Plant Cell Environ. 41, 2250–2262. 10.1111/pce.13205.

96. Maestro, M.C., and Alvarez, J. (1988). The effects of temperature on pollination and pollen tube growth in muskmelon (*Cucumis melo* L.). Sci. Hortic. (Amsterdam) 36, 173–181. 10.1016/0304-4238(88)90051-9.

97. Torabinejad, J., Caldwell, M., Flint, S., and Durham, S. (1998). Susceptibility of pollen to UV-B radiation: an assay of 34 taxa. Am. J. Bot. 85, 360. 10.2307/2446329.

98. Franchi, G.G., Nepi, M., and Pacini, E. (2014). Is flower/corolla closure linked to decrease in viability of desiccation-sensitive pollen? Facts and hypotheses: a review of current literature with the support of some new experimental data. Osterr. Bot. Z. 300, 577–584. 10.1007/s00606-013-0911-x.

99. Cun, S., Zhang, C., Chen, J., Qian, L., Sun, H., and Song, B. (2024). Effects of UV-B radiation on pollen germination and tube growth: A global meta-analysis. Sci. Total Environ. 915, 170097. 10.1016/j.scitotenv.2024.170097.

100. Shivanna, K.R. (2019). Pollen Biology and Biotechnology (CRC Press) 10.1201/9780429187704.

101. Gay, G., Kerhoas, C., and Dumas, C. (1987). Quality of a stress-sensitive *Cucurbita pepo* L. pollen. Planta 171, 82–87. 10.1007/BF00395070.

102. Pacini, E. (1997). Pollen viability related to type of pollination in six angiosperm species. Ann. Bot. 80, 83–87. 10.1006/anbo.1997.0421.

103. Nepi, M. (1993). Pollination, Pollen Viability and Pistil Receptivity in *Cucurbita pepo*. Ann. Bot. 72, 527–536. 10.1006/anbo.1993.1141.

104. Prokop, P., Jersáková, J., Fančovičová, J., and Pipíška, M. (2019). Flower closure enhances pollen viability in *Crocus discolor* G. Reuss. Flora 250, 68–71. 10.1016/j.flora.2018.11.019.

105. Brochu, K.K., van Dyke, M.T., Milano, N.J., Petersen, J.D., McArt, S.H., Nault, B.A., Kessler, A., and Danforth, B.N. (2020). Pollen defenses negatively impact foraging and fitness in a generalist bee (*Bombus impatiens*: Apidae). Sci. Rep. 10, 3112. 10.1038/s41598-020-58274-2.

106. Willis Chan, D.S., and Raine, N.E. (2021). Phenological synchrony between the hoary squash bee (*Eucera pruinosa*) and cultivated acorn squash (*Cucurbita pepo*) flowering is imperfect at a northern site. Curr. Res. Insect Sci. 1, 100022. 10.1016/j.cris.2021.100022.

107. Giurfa, M., and Núñez, J.A. (1992). Honeybees mark with scent and reject recently visited flowers. Oecologia 89, 113–117. 10.1007/BF00319022.

108. Stout, J.C., Goulson, D., and Allen, J.A. (1998). Repellent scent-marking of flowers by a guild of foraging bumblebees (*Bombus* spp.). Behav. Ecol. Sociobiol. 43, 317–326. 10.1007/s002650050497.

109. Witjes, S., and Eltz, T. (2009). Hydrocarbon footprints as a record of bumblebee flower visitation. J. Chem. Ecol. 35, 1320–1325. 10.1007/s10886-009-9720-7.

110. Clarke, D., Whitney, H., Sutton, G., and Robert, D. (2013). Detection and learning of floral electric fields by bumblebees. Science 340, 66–69. 10.1126/science.1230883.

111. Harrap, M.J.M., Hempel de Ibarra, N., Knowles, H.D., Whitney, H.M., and Rands, S.A. (2021). Bumblebees can detect floral humidity. J. Exp. Biol. 224. 10.1242/jeb.240861.

112. Harrison, A.S., and Rands, S.A. (2022). The ability of bumblebees *Bombus terrestris* (Hymenoptera: Apidae) to detect floral humidity is dependent upon environmental humidity. Environ. Entomol. 51, 1010–1019. 10.1093/ee/nvac049.

113. Barman, M., Tenhaken, R., and Dötterl, S. (2024). Negative and sex-specific effects of drought on flower production, resources and pollinator visitation, but not on floral scent in monoecious *Cucurbita pepo*. New Phytol. 244, 1013–1023. 10.1111/nph.20016.

114. Nepi, M., Guarnieri, M., and Pacini, E. (2001). Nectar secretion, reabsorption, and sugar composition in male and female flowers of *Cucurbita pepo*. Int. J. Plant Sci. 162, 353–358. 10.1086/319581.

115. Nepi, M., Cresti, L., Guarnieri, M., and Pacini, E. (2011). Dynamics of nectar production and nectar homeostasis in male flowers of *Cucurbita pepo* L. Int. J. Plant Sci. 172, 183–190. 10.1086/657648.

116. Edge, A.A., van Nest, B.N., Johnson, J.N., Miller, S.N., Naeger, N., Boyd, S.D., and Moore, D. (2012). Diel nectar secretion rhythm in squash (*Cucurbita pepo*) and its relation with pollinator activity. Apidologie (Celle) 43, 1–16. 10.1007/s13592-011-0087-8.

117. Chown, S.L., Sørensen, J.G., and Terblanche, J.S. (2011). Water loss in insects: an environmental change perspective. J. Insect Physiol. 57, 1070–1084. 10.1016/j.jinsphys.2011.05.004.

118. O’Donnell, M.J. (2022). A perspective on insect water balance. J. Exp. Biol. 225. 10.1242/jeb.242358.

119. Benoit, J.B., McCluney, K.E., DeGennaro, M.J., and Dow, J.A.T. (2023). Dehydration dynamics in terrestrial arthropods: From water sensing to trophic interactions. Annu. Rev. Entomol. 68, 129–149. 10.1146/annurev-ento-120120-091609.

120. Davis, C.C., Endress, P.K., and Baum, D.A. (2008). The evolution of floral gigantism. Curr. Opin. Plant Biol. 11, 49–57. 10.1016/j.pbi.2007.11.003.

121. Johnson, S.D. (2016). Carrion flowers. Curr. Biol. 26, R556–R558. 10.1016/j.cub.2015.07.047.

122. Seymour, R.S., White, C.R., and Gibernau, M. (2009). Endothermy of dynastine scarab beetles (*Cyclocephala colasi*) associated with pollination biology of a thermogenic arum lily (*Philodendron solimoesense*). J. Exp. Biol. 212, 2960–2968. 10.1242/jeb.032763.

123. Salzman, S., Dahake, A., Kandalaft, W., Valencia-Montoya, W.A., Calonje, M., Specht, C.D., and Raguso, R.A. (2023). Cone humidity is a strong attractant in an obligate cycad pollination system. Curr. Biol. 33, 1654–1664.e4. 10.1016/j.cub.2023.03.021.

124. Oelschlägel, B., Nuss, M., von Tschirnhaus, M., Pätzold, C., Neinhuis, C., Dötterl, S., and Wanke, S. (2015). The betrayed thief - the extraordinary strategy of *Aristolochia rotunda* to deceive its pollinators. New Phytol. 206, 342–351. 10.1111/nph.13210.

125. Rupp, T., Oelschlägel, B., Rabitsch, K., Mahfoud, H., Wenke, T., Disney, R.H.L., Neinhuis, C., Wanke, S., and Dötterl, S. (2021). Flowers of deceptive *Aristolochia microstoma* are pollinated by phorid flies and emit volatiles known from invertebrate carrion. Front. Ecol. Evol. 9. 10.3389/fevo.2021.658441.

126. Oelschlägel, B., Gorb, S., Wanke, S., and Neinhuis, C. (2009). Structure and biomechanics of trapping flower trichomes and their role in the pollination biology of *Aristolochia* plants (Aristolochiaceae). New Phytol. 184, 988–1002. 10.1111/j.1469-8137.2009.03013.x.

127. Wee, S.L., Tan, S.B., and Jürgens, A. (2018). Pollinator specialization in the enigmatic *Rafflesia cantleyi*: A true carrion flower with species-specific and sex-biased blow fly pollinators. Phytochemistry 153, 120–128. 10.1016/j.phytochem.2018.06.005.

128. Ollerton, J., Masinde, S., Meve, U., Picker, M., and Whittington, A. (2009). Fly pollination in *Ceropegia* (Apocynaceae: Asclepiadoideae): biogeographic and phylogenetic perspectives. Ann. Bot. 103, 1501–1514. 10.1093/aob/mcp072.

129. Vereecken, N.J., Dorchin, A., Dafni, A., Hötling, S., Schulz, S., and Watts, S. (2013). A pollinators’ eye view of a shelter mimicry system. Ann. Bot. 111, 1155– 1165. 10.1093/aob/mct081.

130. Sapir, Y., Shmida, A., and Ne’eman, G. (2006). Morning floral heat as a reward to the pollinators of the *Oncocyclus irises*. Oecologia 147, 53–59. 10.1007/s00442-005-0246-6.

131. Willmer, P., and Stone, G. (1997). Temperature and water relations in desert bees. Journal of Thermal Biology 22, 453–465. 10.1016/S0306-4565(97)00064-8.

132. Dai, A., Zhao, T., and Chen, J. (2018). Climate change and drought: A precipitation and evaporation perspective. Curr. Clim. Change Rep. 4, 301–312. 10.1007/s40641-018-0101-6.

133. Yuan, X., Wang, Y., Ji, P., Wu, P., Sheffield, J., and Otkin, J.A. (2023). A global transition to flash droughts under climate change. Science 380, 187–191. 10.1126/science.abn6301.

134. Sinclair, B.J., Saruhashi, S., and Terblanche, J.S. (2024). Integrating water balance mechanisms into predictions of insect responses to climate change. J. Exp. Biol. 227. 10.1242/jeb.247167.

135. Jones, L.J., Miller, D.A., Schilder, R.J., and López-Uribe, M.M. (2024). Body mass, temperature, and pathogen intensity differentially affect critical thermal maxima and their population-level variation in a solitary bee. Ecol. Evol. 14, e10945. 10.1002/ece3.10945.

