## Supplementary material for "Squash flowers as microhabitats: the effects of floral temperature and humidity on pollen viability and visitor behavior": Document S1

**Video S1** *Cucurbita foetidissima* floral chambers occupied by courting, mating, and fighting *Drosophila pseudoobscura* flies

**Video S2** Tens of *Cucurbit* beetles in the floral chambers of *Cucurbita pepo*

**Video S3** Squash flowers as microhabitats for diverse insects

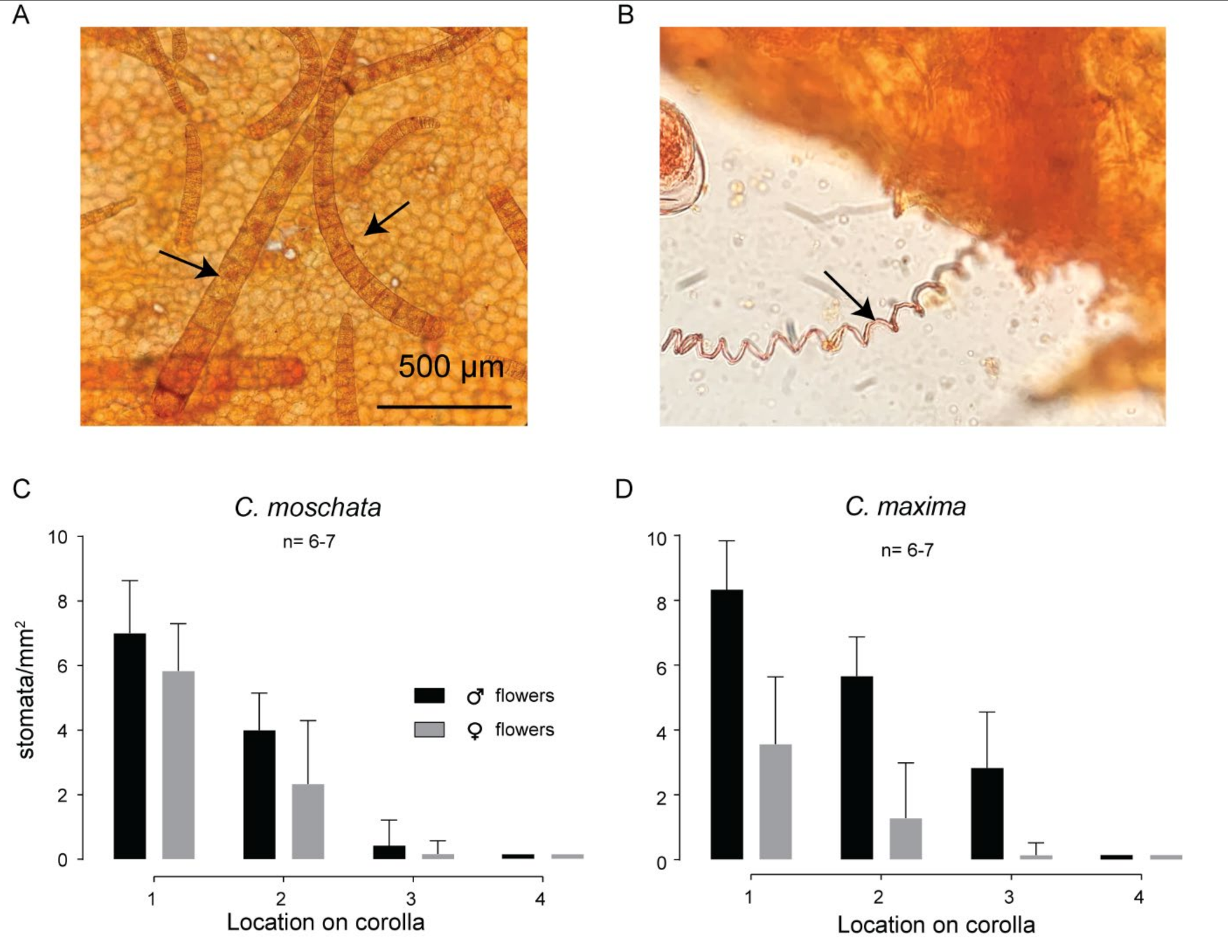


**Fig. S1 Corolla peels showing the different cells on the flower surface including stomatal counts (related to figure 2)**

A. Multicellular trichomes are abundant in the floral chamber on the inner surface of the flower on locations 1 and 2 corresponding to the inset image shown in Fig. 2C. Arrows point toward the trichomes, and the scale bar is shown at the bottom right of the image.
B. Floral peel image shows that flowers are supplied with water through protoxylem (see arrow).
C. And D. Stomatal counts across 4 locations on the inner surface of *Cucurbita moschata* and *Cucurbita maxima* flowers collected from Squash farms near Cornell.

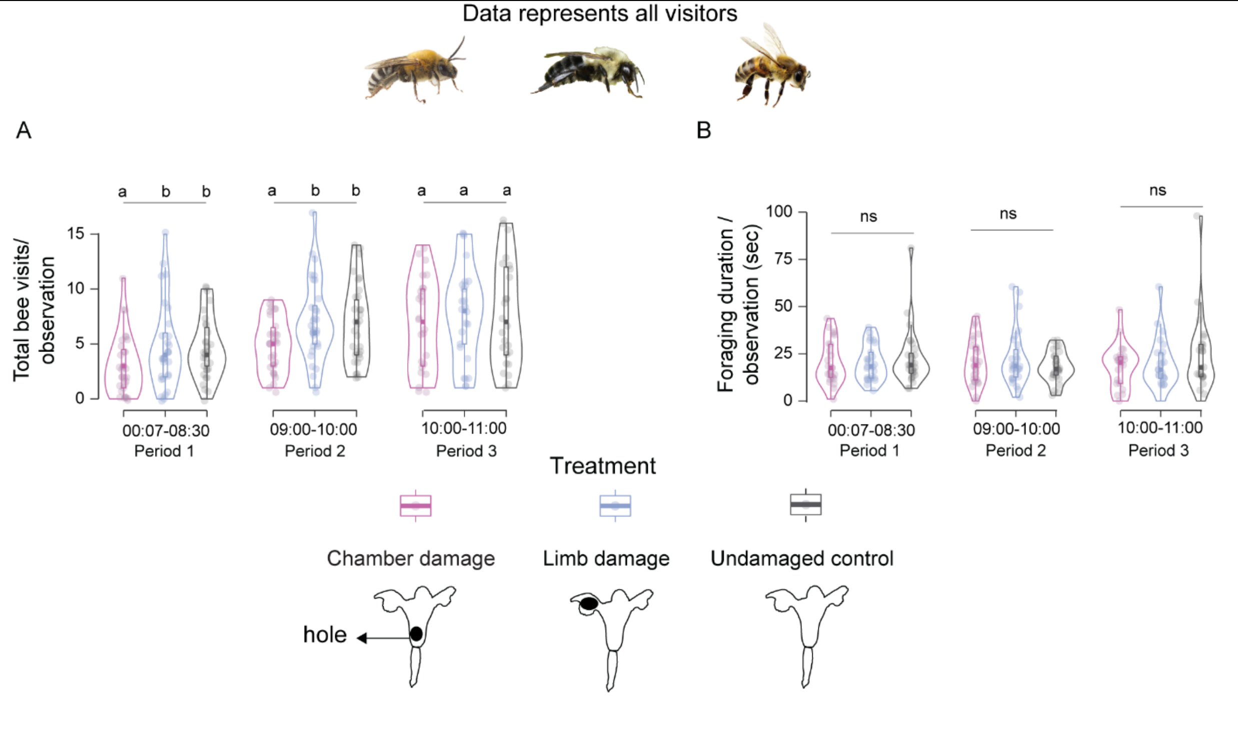


**Fig S2. Effects of humidity-manipulated flowers on the visitation frequency and foraging duration of all bees across three time points (Periods).** (Related to figure 5)

Violin plots with embedded boxplots of bee visitation frequency (a) and foraging time (b) for all bees combined. The data are parsed by period and floral treatment. Colors refer to the treatments shown below. Data are analyzed with a generalized linear model (GLM) and significance is denoted with letters on top of the panel. 'n.s' indicates no significance

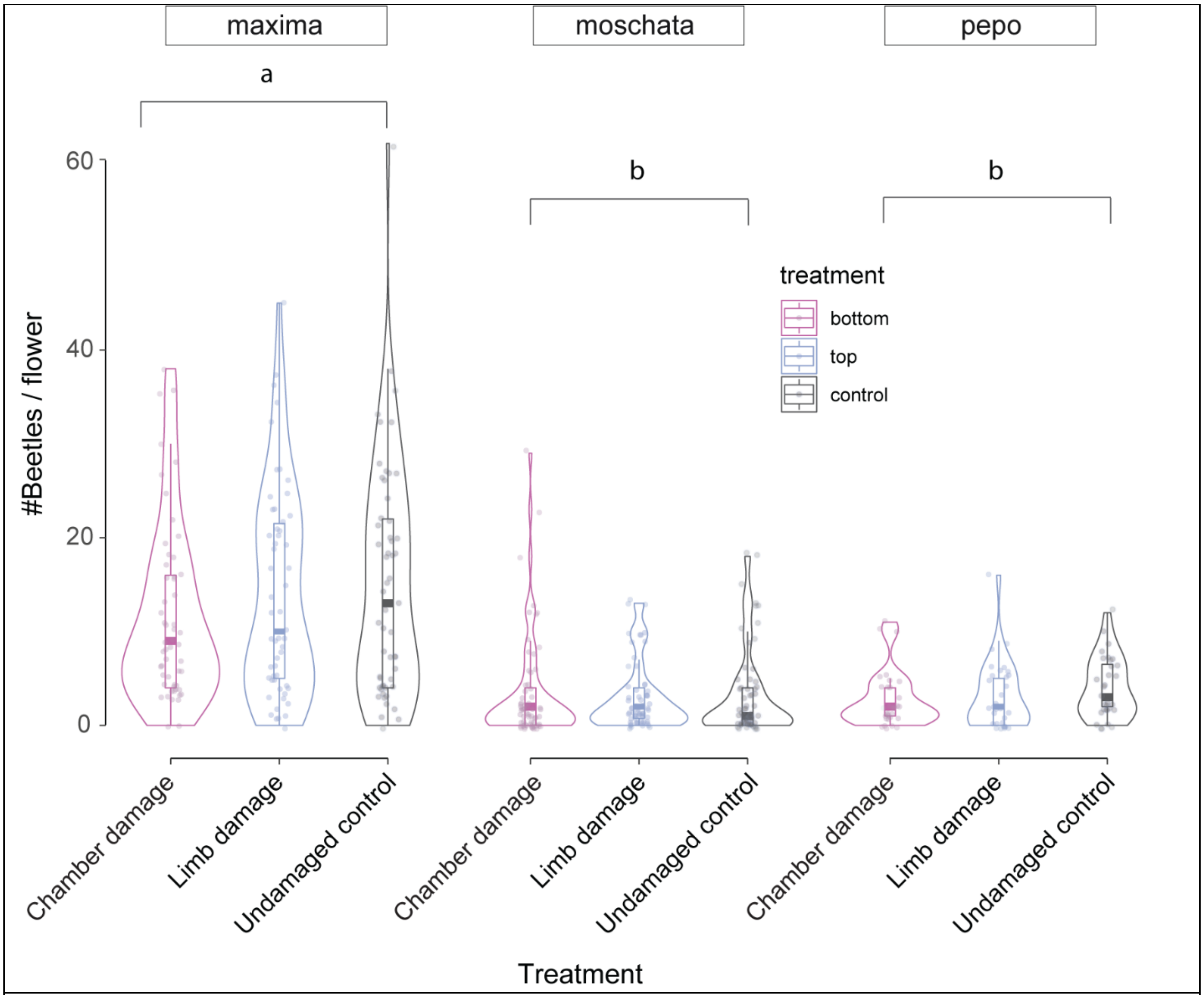


**Fig. S3 Beetle counts within humidity-manipulated flowers separated by plant varieties.** (Related to figure 6)

*C. maxima* flowers attract more beetles than *C. moschata* and *C. pepo* squash varieties. Data are color-coded by treatments and presented as boxplots embedded inside violin plots. Letters on top indicate whether varieties differ from each other (“emmeans” package in R[1]).

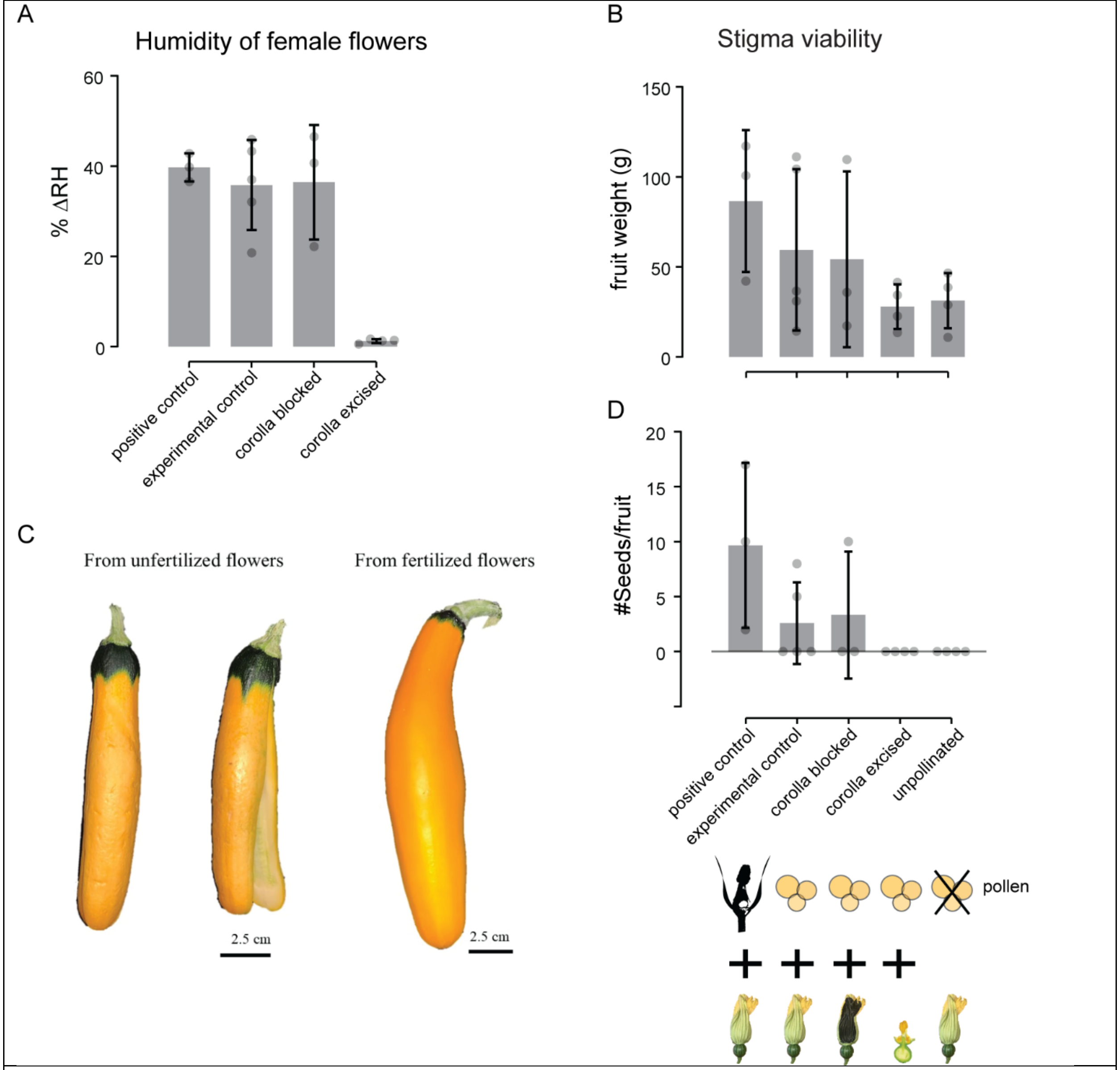


**Fig. S4 Greenhouse experiment on *C. pepo* female flowers to test stigma viability across humidity-manipulated flowers shows an impact on fruit quality and seed set.** (Related to discussion section)
(a) The humidity of the n=3-5 female flowers across experimental treatments. (b) Weights of the golden zucchini shown in (c) across treatments. (d) Number of seeds excavated from dried zucchini. Data are bar plots showing mean ± SD. Dots show individual datapoints. We could not increase sample size for this experiment because the plants succumbed to severe powdery mildew infestation during both experimental trials. The plants typically produced only 1-2 female flowers before they wilted from mildew. Icons respresents the treatments. Individual female flowers underwent one of the following treatments at 8 am: Corolla excised, inside blocked, outside blocked, control, positive control. The flowers were pollinated by hand with outcrossed pollen using a Q-tip. The Q-tip was touched to the anthers and rotated once. This collects ~250 pollen grains per tap, equivalent to a bee visit to the flower [2]. Winsor et al 1987 show that one application of 240 ± 36 produced about 40 seeds. The female flowers were tagged with the treatment, date, and pollination time.

**Table S1. A pairwise comparison between the intercepts of the vertical humidity gradients across treatments.**

Pairwise comparisons of the intercept '*y0*' of floral vertical humidity gradients between squash flowers with different treatments. Parameters are extracted from a nonlinear exponential decay model and compared using 'emmeans' library and p-values adjusted with Tukey-Kramer method

| Flower sex | contrast | estimate | SE | df | t.ratio | P value |
|---|---|---|---|---|---|---|
| **Female** | control – nectar extracted | 1.92 | 0.40 | 11039 | 4.77 | **<0.0001** |
| | control – Petroleum jelly blocked | 10.65 | 0.39 | 11039 | 26.99 | **<0.0001** |
| | Nectar extracted - Petroleum jelly blocked | 8.73 | 0.41 | 11039 | 21.08 | **<0.0001** |
| **Male** | control – nectar extracted | -2.76 | 0.37 | 14287 | -7.46 | **<0.0001** |
| | control - Petroleum jelly blocked | 11.89 | 0.36 | 14287 | 33.41 | **<0.0001** |
| | Nectar extracted - Petroleum jelly blocked | 14.64 | 0.33 | 14287 | 43.79 | **<0.0001** |

**Table S2. A pairwise comparison between the decay rate of the vertical humidity gradients across treatments.**

Pairwise comparisons of the decay rate '*α*' of vertical humidity gradients between squash flowers with different treatments. Parameters are extracted from a nonlinear exponential decay model and compared using 'emmeans' library and p-values adjusted with Tukey-Kramer method

| Flower sex | contrast | estimate | SE | df | t.ratio | P value |
|---|---|---|---|---|---|---|
| **Female** | control – nectar extracted | 0.004 | 0.0005 | 11039 | 8.50 | **<0.0001** |
| | control – Petroleum jelly blocked | -0.007 | 0.0007 | 11039 | -9.58 | **<0.0001** |
| | Nectar extracted - Petroleum jelly blocked | -0.011 | 0.0007 | 11039 | -15.56 | **<0.0001** |
| **Male** | control – nectar extracted | 0.007 | 0.0003 | 14287 | 19.01 | **<0.0001** |
| | control - Petroleum jelly blocked | 0.010 | 0.0003 | 14287 | 27.39 | **<0.0001** |
| | Nectar extracted - Petroleum jelly blocked | 0.003 | 0.0003 | 14287 | 11.52 | **<0.0001** |

**Table S3. Effects of floral treatments on Pollen behavior.**

A two-way ANOVA followed by Tukey's HSD on ΔRH and pollen types encountered across treatments and time. The pollen data is square root transformed. Significant interactions at *P*<0.05 are bolded.

| Time | Df | F | Control-Corolla excised (*P*-value) | Vaseline-Control (*P*-value) | Vaseline-Corolla excised (*P*-value) |
|---|---|---|---|---|---|
| aov(ΔRH~Treatment+Time) | | | | | |
| 07:30 | 2 | 3.63 | **0.0367** | 0.7303 | 0.1696 |

| | | | | | |
|---|---|---|---|---|---|
| 08:30 | 2 | 16.07 | **<.0001** | **0.0030** | 0.1156 |
| 09:30 | 2 | 17.97 | **<.0001** | 0.2894 | **0.0002** |
| 10:30 | 2 | 10.18 | **0.0001** | 0.0545 | 0.0943 |
| 11:30 | 2 | 15.75 | **<.0001** | 0.2694 | **0.0009** |
| 12:30 | 2 | 13.75 | **<.0001** | 0.4377 | **0.0011** |
| aov(sqrt(percent germinated)~Treatment+Time) | | | | | |
| 07:30 | 2 | 0.753 | 0.5251 | 0.9985 | 0.5571 |
| 08:30 | 2 | 1.714 | 0.2183 | 0.9815 | 0.2964 |
| 09:30 | 2 | 12.71 | **<.0001** | 0.3584 | **0.0024** |
| 10:30 | 2 | 11.12 | **<.0001** | 0.4240 | **0.0044** |
| 11:30 | 2 | 6.41 | **0.0030** | 0.6807 | **0.0312** |
| 12:30 | 2 | 11.54 | **<.0001** | 0.1133 | **0.0208** |
| aov(sqrt(percent ruptured)~Treatment+Time) | | | | | |
| 07:30 | 2 | 0.037 | 0.9956 | 0.9616 | 0.9828 |
| 08:30 | 2 | 0.937 | 0.4164 | 0.9787 | 0.5334 |
| 09:30 | 2 | 7.280 | **0.0043** | 0.9997 | **0.0046** |
| 10:30 | 2 | 11.61 | **0.0002** | 0.9866 | **0.0003** |
| 11:30 | 2 | 11.10 | **0.0001** | 0.6591 | **0.0020** |
| 12:30 | 2 | 7.113 | **0.0044** | 0.9961 | **0.0057** |
| aov(sqrt(percent ungerminated)~Treatment+Time) | | | | | |
| 07:30 | 2 | 1.179 | 0.3395 | 0.9799 | 0.4441 |
| 08:30 | 2 | 0.028 | 0.9982 | 0.9716 | 0.9839 |
| 09:30 | 2 | 2.043 | 0.2541 | 0.1559 | 0.9599 |
| 10:30 | 2 | 3.414 | 0.1165 | 0.9041 | **0.0451** |
| 11:30 | 2 | 4.498 | **0.0155** | 0.7842 | 0.0793 |
| 12:30 | 2 | 3.411 | 0.0916 | 0.9691 | 0.0537 |

**Table S4 Model coefficients obtained from Linear mixed effect models on the ΔT and ΔRH of experimentally manipulated flowers in field experiments.**
In these models, ΔT or ΔRH served as the response variable, treatment * period was the fixed effect explanatory variable, and plant variety, day, and site were included as random effects in the initial model and dropped in subsequent models if they did not contribute to the variance of the model. For ΔRH we performed pairwise comparisons between treatments within a period using library ‘emmeans’, Kenward-roger method for degrees of freedom, and Tukey-Kramer adjusted p-values.

| Model | Variable | Estimate | SE | df | t value | Pr (> \| t \|) |
|---|---|---|---|---|---|---|
| Floral ΔT | | | | | | |
| | Intercept | -0.1935 | 0.3772 | 5.3 | -0.513 | 0.629 |
| | Treatment Limb-damaged flowers | 0.0527 | 0.2524 | 237.6 | 0.209 | 0.835 |
| | Treatment control flowers | 0.2795 | 0.2501 | 237.4 | 1.118 | 0.265 |
| | Period 2 | 0.2439 | 0.2515 | 241.5 | 0.970 | 0.333 |
| | Period 3 | 0.0622 | 0.2655 | 239.1 | 0.234 | 0.815 |
| | Treatment Limb-damaged flowers: Period 2 | 0.1761 | 0.3553 | 237.5 | 0.496 | 0.621 |

| | | | | | | |
|---|---|---|---|---|---|---|
| | Treatment Undamaged control flowers: Period 2 | 0.0337 | 0.3537 | 237.4 | 0.095 | 0.924 |
| | Treatment Limb-damaged flowers: Period 3 | -0.0518 | 0.3758 | 237.5 | -0.138 | 0.890 |
| | Treatment Undamaged control flowers: Period 3 | -0.0679 | 0.3743 | 237.4 | -0.181 | 0.856 |
| Floral ΔRH | | | | | | |
| | Intercept | 4.4906 | 2.0422 | 3.8 | 2.199 | 0.095 |
| | Treatment Limb-damaged flowers | 6.7357 | 0.6785 | 238.7 | 9.927 | **<0.0001** |
| | Treatment Undamaged control flowers | 5.9571 | 0.6764 | 238.6 | 8.808 | **<0.0001** |
| | Period 2 | 2.9837 | 0.6764 | 244.2 | 4.411 | **<0.0001** |
| | Period 3 | 7.4133 | 0.7037 | 240.2 | 10.535 | **<0.0001** |

**Table S5. Effects of floral humidity manipulations on the visitation frequency of insects.** Model coefficients obtained from GLMMs with Poisson error distribution on the frequency of bee visits to flowers of different treatments across three periods. Counts of individual bees served as the response variable, treatment * period was the fixed effect and plant variety, day, and site were included as random effects in the initial model and dropped in subsequent models if they did not contribute to the variance of the model. For honeybee visitation frequency we used a zero-inflated GLM with Poisson log link due to the fewer counts of honeybees. For GLMs where the treatment had a significant impact on visitation rates, they were compared pairwise within a period using library (emmeans) with Tukey-Kramer adjusted p-values.

| Model | Variable | Estimate | SE | *z* value | Pr (> \| *z* \|) |
|---|---|---|---|---|---|
| All bee species visitation frequency | | | | | |
| | Intercept | 1.1165 | 0.1890 | 5.90 | **<0.0001** |
| | Limb-damaged flowers | 0.4129 | 0.1327 | 3.11 | **0.001** |
| | Undamaged-Control flowers | 0.4019 | 0.1323 | 3.03 | **0.002** |
| | Period 2 | 0.4649 | 0.1319 | 3.52 | **< 0.001** |
| | Period 3 | 0.8151 | 0.1281 | 6.35 | **<0.0001** |
| | Treatment Limb-damaged flowers: Period 2 | -0.0437 | 0.1698 | -0.25 | 0.7969 |
| | Treatment Undamaged control flowers: Period 2 | -0.0559 | 0.1698 | -0.33 | 0.7416 |

| | | | | | |
|---|---|---|---|---|---|
| | Treatment Limb-damaged flowers: Period 3 | -0.2868 | 0.1690 | -1.69 | 0.0898 |
| | Treatment Undamaged control flowers: Period 3 | -0.3018 | 0.1691 | -1.78 | 0.0784 |
| Squash bee visitation frequency | | | | | |
| | Intercept | -0.1000 | 0.3665 | -0.27 | 0.7849 |
| | Limb-damaged flowers | 0.6945 | 0.2071 | 3.35 | **0.0008** |
| | Undamaged-Control flowers | 0.6343 | 0.2081 | 3.04 | **0.0023** |
| | Period 2 | 0.4382 | 0.2185 | 2.00 | **0.0449** |
| | Period 3 | 0.7961 | 0.2157 | 3.69 | **0.0002** |
| | Treatment Limb-damaged flowers: Period 2 | -0.0307 | 0.2678 | -0.11 | 0.9086 |
| | Treatment Undamaged control flowers: Period 2 | -0.2288 | 0.2738 | -0.83 | 0.4034 |
| | Treatment Limb-damaged flowers: Period 3 | -0.6423 | 0.2781 | -2.30 | **0.0209** |
| | Treatment Undamaged control flowers: Period 3 | -0.5487 | 0.2778 | -1.97 | **0.0482** |
| Bumblebee visitation frequency | | | | | |
| | Intercept | 0.4057 | 0.2756 | 1.47 | 0.1410 |
| | Limb-damaged flowers | 0.1716 | 0.1832 | 0.93 | 0.3490 |
| | Undamaged-Control flowers | 0.1035 | 0.1852 | 0.55 | 0.5763 |
| | Period 2 | 0.4270 | 0.1748 | 2.44 | **0.0146** |
| | Period 3 | 0.7356 | 0.1676 | 4.33 | **<0.0001** |
| | Treatment Limb-damaged flowers: Period 2 | -0.0191 | 0.2351 | -0.08 | 0.9350 |
| | Treatment Undamaged control flowers: Period 2 | 0.1542 | 0.2345 | 0.65 | 0.5107 |
| | Treatment Limb-damaged flowers: Period 3 | -0.0763 | 0.2292 | -0.33 | 0.7392 |
| | Treatment Undamaged control flowers: Period 3 | -0.0547 | 0.2317 | -0.23 | 0.8133 |
| Honeybee visitation frequency | | | | | |

| | | | | | |
|---|---|---|---|---|---|
| | Intercept | -0.1345 | 0.9439 | -0.14 | 0.8866 |
| | Limb-damaged flowers | -1.8801 | 1.0685 | -1.76 | 0.0785 |
| | Undamaged-Control flowers | 0.4699 | 1.0163 | 0.46 | 0.6438 |
| | Period 2 | 0.0001 | 1.1559 | 0.00 | 0.9999 |
| | Period 3 | 0.1613 | 1.0879 | 0.14 | 0.8821 |
| | Treatment Limb-damaged flowers: Period 2 | 2.0412 | 1.3710 | 1.48 | 0.1365 |
| | Treatment Undamaged control flowers: Period 2 | -0.3087 | 1.2746 | -0.24 | 0.8086 |
| | Treatment Limb-damaged flowers: Period 3 | 1.8123 | 1.2692 | 1.42 | 0.1533 |
| | Treatment Undamaged control flowers: Period 3 | -0.0307 | 1.2223 | -0.02 | 0.9799 |

**Table S6. Effects of floral humidity manipulations on the foraging duration of insects.** Model coefficients obtained from LMMs on the average time spent by bees in the flowers with the different treatments across three periods. Foraging time of individual bees served as the response variable, treatment * period was the fixed effect, and plant variety, day, and site were included as random effects in the initial model and dropped in subsequent models if they did not contribute to the variance of the model.

| Model | Variable | Estimate | SE | *df* | *t* value | Pr (> \| *t* \|) |
|---|---|---|---|---|---|---|
| All bee species' foraging duration | | | | | | |
| | Intercept | 19.259 | 10.180 | 121.045 | 1.892 | 0.0609 |
| | Limb-damaged flowers | 4.106 | 13.594 | 231.110 | 0.302 | 0.7629 |
| | Undamaged-Control flowers | 14.154 | 13.156 | 232.306 | 1.076 | 0.2831 |
| | Period 2 | 28.031 | 13.110 | 237.215 | 2.138 | **0.0335** |
| | Period 3 | 7.219 | 13.780 | 235.746 | 0.524 | 0.6009 |
| | Treatment Limb-damaged flowers: Period 2 | -17.189 | 18.350 | 231.044 | -0.937 | 0.3499 |
| | Treatment Undamaged control flowers: Period 2 | -29.997 | 18.027 | 231.690 | -1.664 | 0.0975 |
| | Treatment Limb-damaged flowers: Period 3 | -12.033 | 19.317 | 231.036 | -0.623 | 0.5340 |

| | | | | | | |
|---|---|---|---|---|---|---|
| | Treatment Undamaged control flowers: Period 3 | -11.887 | 19.012 | 231.618 | -0.625 | 0.5324 |
| Squash bee foraging duration | | | | | | |
| | Intercept | 24.210 | 11.160 | 7.219 | 2.169 | 0.0655 |
| | Limb-damaged flowers | 1.732 | 11.255 | 173.762 | 0.154 | 0.8778 |
| | Undamaged-Control flowers | 13.606 | 10.786 | 173.269 | 1.261 | 0.2088 |
| | Period 2 | 0.193 | 11.258 | 173.458 | 0.017 | 0.9863 |
| | Period 3 | 4.810 | 11.564 | 173.285 | 0.416 | 0.6779 |
| | Treatment Limb-damaged flowers: Period 2 | 14.422 | 15.346 | 173.474 | 0.940 | 0.3486 |
| | Treatment Undamaged control flowers: Period 2 | -22.987 | 14.950 | 173.105 | -1.538 | 0.1260 |
| | Treatment Limb-damaged flowers: Period 3 | -11.643 | 16.489 | 173.262 | -0.706 | 0.4811 |
| | Treatment Undamaged control flowers: Period 3 | -17.584 | 16.186 | 173.032 | -1.086 | 0.2788 |
| bumblebee foraging duration | | | | | | |
| | Intercept | 23.692 | 17.100 | 17.219 | 1.385 | 0.1836 |
| | Limb-damaged flowers | -1.291 | 18.411 | 191.963 | -0.070 | 0.9442 |
| | Undamaged-Control flowers | 9.337 | 18.547 | 191.046 | 0.503 | 0.6153 |
| | Period 2 | 46.048 | 17.997 | 194.523 | 2.559 | **0.0113** |
| | Period 3 | -5.265 | 18.305 | 193.703 | -0.288 | 0.7740 |
| | Treatment Limb-damaged flowers: Period 2 | -44.896 | 23.835 | 190.358 | -1.884 | 0.0611 |
| | Treatment Undamaged control flowers: Period 2 | -53.861 | 23.676 | 190.125 | -2.275 | **0.0240** |
| | Treatment Limb-damaged flowers: Period 3 | 3.527 | 24.168 | 191.184 | 0.146 | 0.8841 |
| | Treatment Undamaged control flowers: Period 3 | -5.391 | 24.280 | 190.219 | -0.222 | 0.8245 |
| Honeybee foraging duration | | | | | | |

| | | | | | | |
|---|---|---|---|---|---|---|
| | Intercept | 23.771 | 62.515 | 39.341 | 0.380 | 0.706 |
| | Limb-damaged flowers | -18.359 | 76.309 | 41.996 | -0.241 | 0.811 |
| | Undamaged-Control flowers | -8.842 | 70.570 | 41.839 | -0.125 | 0.901 |
| | Period 2 | -17.290 | 76.367 | 41.908 | 0.226 | 0.822 |
| | Period 3 | 3.892 | 73.529 | 41.010 | 0.053 | 0.958 |
| | Treatment Limb-damaged flowers: Period 2 | 145.558 | 96.587 | 41.911 | 1.507 | 0.139 |
| | Treatment Undamaged control flowers: Period 2 | 51.938 | 87.795 | 42.000 | 0.592 | 0.557 |
| | Treatment Limb-damaged flowers: Period 3 | 15.330 | 90.652 | 41.782 | 0.169 | 0.867 |
| | Treatment Undamaged control flowers: Period 3 | 83.929 | 89.761 | 41.133 | 0.935 | 0.355 |

**Table S7. Effects of floral humidity manipulations on the residence of beetles and squash bees.** Model coefficients obtained from a GLM with Poisson distribution on the occurrence of cucurbit beetles residing inside flowers with different experimental treatments. For squash bee occurrence in flowers, we used a zero-inflated GLM with Poisson distribution.

| Model | Variable | Estimate | SE | *z* value | Pr (> \| *z* \|) |
|---|---|---|---|---|---|
| Cucurbit beetles | | | | | |
| | Intercept | 2.8484 | 0.6339 | 4.493 | **<0.0001** |
| | Treatment Limb-damaged flowers | 0.0989 | 0.0447 | 2.213 | **0.0269** |
| | Treatment Undamaged-control flowers | 0.1993 | 0.0436 | 4.563 | **<0.0001** |
| | Variety moschata | -1.5131 | 0.2769 | -5.464 | **<0.0001** |
| | Variety pepo | -1.4060 | 0.0609 | -23.076 | **<0.0001** |
| | Site | -0.7667 | 0.6696 | -1.145 | 0.2522 |
| Squash bees | | | | | |
| | Intercept | 0.1828 | 1.1467 | 0.159 | 0.8733 |
| | Treatment Limb-damaged flowers | 1.1457 | 0.3960 | 2.893 | **0.0038** |
| | Treatment Undamaged-control flowers | 1.0086 | 0.4013 | 2.513 | **0.0119** |
| | Variety moschata | -0.8125 | 1.0395 | -0.782 | 0.4344 |
| | Variety pepo | -0.5450 | 0.3961 | -1.376 | 0.1688 |
| | Site | -1.3613 | 1.0499 | -1.297 | 0.1947 |

**Table S8**. **Squash flower morphometrics**. Values show mean ± SD rounded up to 1 decimal point and numbers in parentheses indicate sample sizes. We were not able to collect data on floral dry mass and % water weight for *C. foetidissima* because we sacrificed those flowers for stomatal counts.

| Species | Flower Sex | Corolla diameter (cm) | Corolla Height from nectary (cm) | Fresh mass (g) | Dry mass (g) | % Water weight | Flower chamber volume ($cm^3$) |
|---|---|---|---|---|---|---|---|
| *Cucurbita foetidissima* | male | 6.6 ± 1.4 (8) | 9.0 ± 0.7 (8) | 7.0 ± 2.1 (7) | NA | NA | 41.0 ± 24.5 (7) |
| | female | 7.6 ± 2.9 (8) | 9.3 ± 1.2 (8) | 12.0 ± 2.3 (7) | NA | NA | 38.4 ± 15.9 (7) |
| *Cucurbita pepo* | male | 11.6 ± 1.0 (14) | 8.8 ± 1.1 (13) | 4.13 ± 0.6 (10) | 0.3 ± 0.04 (10) | 92.92 ± 0.9 (10) | 29.3 ± 7.3 (10) |
| | female | 13.1 ± 2.4 (5) | 8.9 ± 1.0 (5) | 14.8 ± 5.1 (5) | 0.9 ± 0.4 (5) | 94.0 ± 0.8 (5) | 41.9 ± 19.8 (5) |
| *Cucurbita moschata* | male | 12.4 ± 1.3 (13) | 8.5 ± 1.0 (13) | 5.9 ± 1.3 (7) | 0.3 ± 0.1 (7) | 95.0 ± 0.4 (7) | 29.0 ± 6.4 (7) |
| | female | 11.0 ± 1.8 (10) | 7.4 ± 0.9 (10) | 13.2 ± 5.1 (10) | 0.7 ± 0.3 (10) | 94.6 ± 0.3 (10) | 24.3 ± 9.1 (10) |
| *Cucurbita maxima* | male | 11.9 ± 2.2 (12) | 8.9 ± 1.2 (12) | 7.3 ± 1.8 (6) | 0.4 ± 0.1 (6) | 94.1 ± 0.6 (6) | 31.8 ± 5.3 (6) |
| | female | 14.2 ± 2.7 (10) | 10.1 ± 1.2 (10) | 18.2 ± 6.3 (10) | 0.9 ± 0.3 (10) | 94.8 ± 0.4 (10) | 51.6 ± 18.0 (6) |

### References


1. Lenth, R.V. (2024). emmeans: Estimated Marginal Means, aka Least-Squares Means.

2. Winsor, J.A., Davis, L.E., and Stephenson, A.G. (1987). The relationship between pollen load and fruit maturation and the effect of pollen load on offspring vigor in *Cucurbita pepo*. Am. Nat. *129*, 643–656.
